# sdmTMB: An R Package for Fast, Flexible, and User-Friendly Generalized Linear Mixed Effects Models with Spatial and Spatiotemporal Random Fields

**DOI:** 10.1101/2022.03.24.485545

**Authors:** Sean C. Anderson, Eric J. Ward, Philina A. English, Lewis A. K. Barnett, James T. Thorson

## Abstract

Geostatistical spatial or spatiotemporal data are common across scientific fields. However, appropriate models to analyse these data, such as generalised linear mixed effects models (GLMMs) with Gaussian Markov random fields (GMRFs), are computationally intensive and challenging for many users to implement. Here, we introduce the R package **sdmTMB**, which extends the flexible interface familiar to users of **lme4, glmmTMB**, and **mgcv** to include spatial and spatiotemporal latent GMRFs using an SPDE-(stochastic partial differential equation) based approach. SPDE matrices are constructed with **fmesher** and estimation is conducted via maximum marginal likelihood with **TMB** or via Bayesian inference with **tmbstan** and **rstan**. We describe the model and explore case studies that illustrate **sdmTMB**’s flexibility in implementing penalised smoothers, non-stationary processes (time-varying and spatially varying coefficients), hurdle models, cross-validation and anisotropy (directionally dependent spatial correlation). Finally, we compare the functionality, speed, and interfaces of related software, demonstrating that **sdmTMB** can be an order of magnitude faster than R-**INLA**. We hope **sdmTMB** will help open this useful class of models to a wider field of geostatistical analysts.

## 1. Introduction

Data are often collected at particular locations in space or in space repeatedly over time. While such data are a rich source of information across many fields, they are challenging to properly model—data closer in space and time are usually more similar to each other than data farther apart due to measured and unmeasured variables (Cressie 1993; Diggle and Ribeiro 2007; Cressie and Wikle 2011). While measured variables can be accounted for with predictors in a model (e.g., measuring and modelling temperature effects on species abundance), unmeasured variables can cause residual spatial correlation. Accounting for this residual correlation is important because doing so allows for valid statistical inference (Legendre and Fortin 1989; Dormann, McPherson, Araújo, Bivand, Bolliger, Carl, Davies, Hirzel, Jetz, Kissling, Kühn, Ohlemüller, Peres-Neto, Reineking, Schröder, Schurr, and Wilson 2007), can improve predictions (e.g., Shelton, Thorson, Ward, and Feist 2014), and can provide useful spatial summary statistics (e.g., Thorson 2019b; Barnett, Ward, and Anderson 2021).

Geostatistical generalized linear mixed effects models (GLMMs) with spatially correlated random effects constitute an appropriate class of models for such data (Rue and Held 2005; Diggle and Ribeiro 2007; Cressie and Wikle 2011; Thorson and Kristensen 2024). Similarly to how random intercepts can account for correlation among groups, spatial or spatiotemporal random effects can account for unmeasured variables that cause observations to be correlated in space or both space and time. A common approach to modelling these spatial effects is with Gaussian random fields (GRFs), where the random effects describing the spatial patterning are assumed to be drawn from a multivariate normal distribution, constrained by covariance functions such as the exponential or Matérn (Cressie 1993; Chilés and Delfiner 1999; Diggle and Ribeiro 2007).

Such models quickly become computationally challenging due to the need to invert large matrices to account for covariation when evaluating the multivariate normal density function. Many solutions have been proposed, such as predictive processes (Banerjee, Gelfand, Finley, and Sang 2008; Latimer, Banerjee, Sang Jr, Mosher, and Silander Jr 2009), the stochastic partial differential equation (SPDE) approximation to GRFs (Lindgren, Rue, and Lindström 2011), and nearest-neighbour Gaussian processes (Datta, Banerjee, Finley, and Gelfand 2016; Finley, Datta, and Banerjee 2022). These approaches aim to minimize the scale of the co-variance estimation problem while providing a means to evaluate the data likelihood, thereby allowing fitting via Bayesian (Gelfand and Banerjee 2017) or maximum likelihood methods. This can greatly improve computational efficiency (e.g., Heaton, Datta, Finley, Furrer, Guinness, Guhaniyogi, Gerber, Gramacy, Hammerling, Katzfuss, Lindgren, Nychka, Sun, and Zammit-Mangion 2019). The SPDE approach has been widely adopted, especially via the R-**INLA** package (Rue, Martino, and Chopin 2009; Lindgren *et al*. 2011; Lindgren and Rue 2015) and an implementation in **TMB** (Template Model Builder, Kristensen, Nielsen, Berg, Skaug, and Bell 2016) that relies on R-**INLA** to create input matrices (Thorson, Skaug, Kristensen, Shelton, Ward, Harms, and Benante 2015c; Thorson 2019a; Osgood-Zimmerman and Wakefield 2023; Thorson and Kristensen 2024). This SPDE method approximates a Matérn correlation function as arising mechanistically from local diffusion in space and/or time (Lindgren *et al*. 2011) and it results in a sparse precision matrix that permits efficient computation using existing sparse-matrix libraries (Rue and Held 2005).

Software packages designed for specifying statistical models that incorporate the SPDE, such as R-**INLA** and **TMB**, are flexible and powerful but can be challenging for many applied researchers. For example, **TMB** requires the user to program in a C++ template and it can be slow to experiment with multiple models when writing bespoke model code. While packages such as **lme4** (Bates, Mächler, Bolker, and Walker 2015) and **glmmTMB** (Brooks, Kristensen, van Benthem, Magnusson, Berg, Nielsen, Skaug, Maechler, and Bolker 2017) let users quickly iterate and explore statistical models—focusing on evaluating fit and comparing models—they lack built-in SPDE or spatiotemporal functionality. Packages such as **VAST** (Thorson 2019a), **tinyVAST** (Thorson, Anderson, Goddard, and Rooper 2024), and **inlabru** (Bachl, Lindgren, Borchers, and Illian 2019) are powerful user interfaces to fit spatial models that use the SPDE, but they either lack a modular interface familiar to those who have used **lme4** or **glmmTMB**, lack some functionality, or may be challenging to learn for some users.

Here, we introduce the R package **sdmTMB**, which implements geostatistical spatial and spatiotemporal GLMMs using **TMB** for model fitting and **fmesher** or R-**INLA** to set up SPDE matrices. Our aim is not to replace the above-mentioned statistical packages, but to provide a fast, flexible, and user-friendly interface that is familiar to users of **lme4, glmmTMB**, or **mgcv** (Wood 2017) for a specific class of spatial and spatiotemporal models. Many individual features of **sdmTMB** may be found in other software (Table 1), but to date this full suite of useful features has not been integrated into a single package. One common application in the field of ecology is species distribution models (SDMs), hence the package name (i.e., SDMs with **TMB**), although the package is widely applicable to other fields and any geostatistical data collected continuously in space and approximated in discrete time-intervals.

**Table 1:** Comparison of functionality between several R packages that can fit geostatistical GLMMs. At the time of writing, the feature set of **tinyVAST** is rapidly evolving and so is not shown here. Notes: ^1^Technically possible but non-trivial. ^2^Penalized smoother GAMs that determine ‘wiggliness’. ^3^**inlabru** but not R-INLA. ^4^Spatiotemporal fields as random walk only. ^5^SPDE approach as in Miller *et al*. (2019). ^6^Does have predictive process knots. ^7^Zero-inflated NB2 only. ^8^Tweedie power parameter fixed for `mgcv::gamm()`. ^9^Possible as log-linked Poisson GLMM with aggregated data. ^10^Hurdle models possible by fitting components separately.

|  | sdmTMB | VAST | R-INLA/inlabru | mgcv | spBayes | spaMM |
| --- | --- | --- | --- | --- | --- | --- |
| Time-varying coefficients | ✓ | —<sup>1</sup> | ✓ | ✓ | ✓ | — |
| Spatially varying coefficients (SVC) | ✓ | ✓ | ✓ | ✓ | ✓ | — |
| GAMs<sup>2</sup> | ✓ | — | ✓ | ✓ | — | — |
| Threshold covariates | ✓ | — | ✓<sup>3</sup> | — | — | — |
| Offsets | ✓ | ✓ | ✓ | ✓ | ✓ | ✓ |
| Spatiotemporal fields | ✓ | ✓ | ✓ | ✓ | ✓<sup>4</sup> | — |
| Spatial + spatiotemporal fields | ✓ | ✓ | ✓ | ✓ | — | — |
| Anisotropy | ✓ | ✓ | — | ✓ | — | — |
| Correlation barriers | ✓ | ✓ | ✓ | ✓ | — | — |
| Separate range parameters for fields | ✓ | — | ✓ | ✓ | — | — |
| Share range parameters across fields | ✓ | ✓ | ✓ | — | — | — |
| SPDE-based | ✓ | ✓ | ✓ | ✓<sup>5</sup> | —<sup>6</sup> | ✓ |
| NB1 distribution | ✓ | — | ✓ | ✓ | — | ✓ |
| NB2 distribution | ✓ | ✓<sup>7</sup> | ✓ | ✓ | — | ✓ |
| Zero-truncated distributions | ✓ | — | ✓ | — | — | ✓ |
| Zero-inflated distributions | ✓ | ✓ | ✓ | — | — | ✓ |
| Tweedie distribution | ✓ | ✓ | ✓ | ✓<sup>8</sup> | — | — |
| Student-t distribution | ✓ | — | ✓ | ✓ | — | — |
| Censored Poisson distribution | ✓ | — | ✓ | — | — | — |
| log Gaussian Cox processes | —<sup>9</sup> | —<sup>9</sup> | ✓ | —<sup>9</sup> | —<sup>9</sup> | —<sup>9</sup> |
| Multivariate responses | — | ✓ | ✓ | ✓ | ✓ | ✓ |
| Built-in delta/hurdle models | ✓ | ✓ | ✓ | —<sup>10</sup> | — | ✓ |
| Poisson-link delta models | ✓ | ✓ | ✓ | — | — | — |
| Likelihood weights | ✓ | — | ✓ | ✓ | ✓ | ✓ |
| Maximum/marginal likelihood | ✓ | ✓ | — | ✓ | — | — |
| Bayesian/optionally Bayesian | ✓ | ✓ | ✓ | ✓ | ✓ | — |
| Priors/penalties | ✓ | — | ✓ | — | ✓ | — |
| Matern PC priors | ✓ | — | ✓ | — | — | — |
| Spatial (or spatial dynamic) factor analysis | — | ✓ | — | — | — | — |
| Empirical Orthogonal Function (EOF) analysis | — | ✓ | — | — | — | — |
| Built-in area-weighted index standardization | ✓ | ✓ | — | — | — | — |
| Built-in cross-validation | ✓ | — | — | — | — | — |

This paper describes the statistical models underlying **sdmTMB** (Section 2), explains how **sdmTMB** is designed and summarizes its software functionality (Section 3), illustrates its use through three case studies (Sections 4, 5, and 6), compares **sdmTMB** to other packages (Section 7) and concludes with a discussion of links to other packages and future development directions (Section 8).

## 2. Model description

### 2.1. A spatial Gaussian random field GLMM

A GLMM with spatial Gaussian random fields can be written as

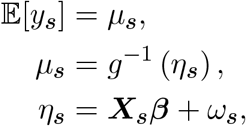

where the expected value 𝔼 [.] of an observation *y* at coordinates in space ***s*** is defined as the mean *µ*_***s***_. That mean *µ*_***s***_ is the result of an inverse link function *g*^−1^ applied to a linear predictor *η*_***s***_. In this case, that linear predictor is the combination of a model matrix ***X***_***s***_ multiplied by a vector of coefficients ***β*** and a value from a spatial random field *ω*_***s***_. This spatial random field represents the effect of latent spatial variables that are not otherwise accounted for in the model. Alternatively, *ω*_***s***_ can be thought of representing spatially correlated noise arising from unmodeled processes. More simply, the vector of *ω*_***s***_ (***ω***) represents a “wiggly” surface with an expected value of zero that is added to the linear predictor in link space (e.g., Figure 1c). The vector ***ω*** is assumed drawn from a multivariate normal distribution with a covariance matrix ∑_*ω*_,

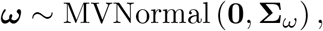

constrained by some function that defines the rate at which spatial covariance decays with distance.

**Figure 1:**
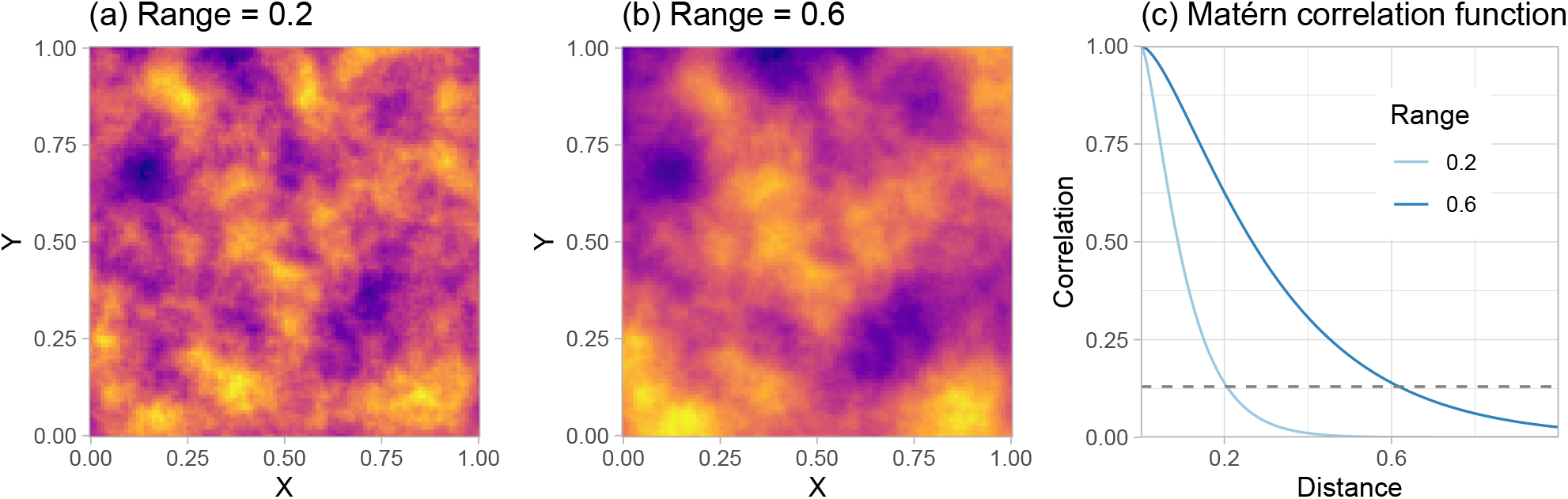
Example Gaussian random fields for two range values. The range describes the distance at which spatial correlation decays to ≈ 0.13 in coordinate units (i.e., the distance at which two points are effectively independent). Panel (a) shows a shorter range than panel (b), which results in a “wigglier” surface. Panel (c) shows the Matérn function for these two range values. The dashed horizontal line shows a correlation of 0.13.

### 2.2. The Matérn covariance function

Various covariance functions are possible, but a popular and flexible one is the Matérn (Whittle 1954; Matérn 1986) (Figure 1). If we define ∥*h*∥ as the Euclidean distance between spatial points ***s***_*j*_ and ***s***_*k*_, we can represent the Matérn covariance Φ as

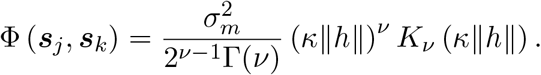

The parameter 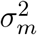 is the marginal variance (magnitude of the random field “wiggles”), Γ represents the Gamma function, *K*_*ν*_ represents the modified Bessel function of the second kind, and *κ* represents the spatial decorrelation rate. The parameter *ν* controls the smoothness of the covariance function. In practice, *ν* is challenging to estimate and herein is fixed at *ν* = 1 (Lindgren *et al*. 2011). A more interpretable derived parameter than the spatial decorrelation rate *κ* is the spatial range—a distance at which two points are effectively independent. A common definition is 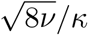 (so if *ν* = 1, range = 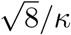), which has the empirically derived property of the distance at which correlation decays to ≈ 0.13 (Lindgren *et al*. 2011) (Figure 1c).

### 2.3. Geometric anisotropy

The assumption that correlation decays equally in all directions can be relaxed to allow for geometric anisotropy

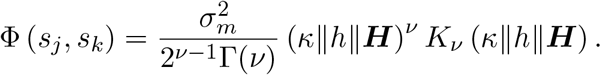

Here, we add a linear transformation matrix ***H*** with two estimated parameters governing the major axis direction of geometric anisotropy and the ratio of the major and minor axes (Haskard 2007; Lindgren *et al*. 2011; Thorson, Shelton, Ward, and Skaug 2015b).

### 2.4. The SPDE approach

In practice, working with the dense covariance matrix ∑ is computationally expensive and methods for working directly with its inverse, the precision matrix ***Q***, are more efficient (***Q*** = ∑^−1^) (Rue and Held 2005; Simpson, Lindgren, and Rue 2012). One such approach is the SPDE approach, which approximates a mechanistic process of local diffusion using methods drawn from finite-element analysis. A full description of the SPDE approach is beyond the scope of this paper. Instead, we refer to the following literature: Lindgren *et al*. (2011) develop the approach. Lindgren and Rue (2015) and Bakka, Rue, Fuglstad, Riebler, Bolin, Illian, Krainski, Simpson, and Lindgren (2018) summarize the SPDE approach for spatial modelling in the context of R-**INLA**. The second chapter of Krainski, Gómez-Rubio, Bakka, Lenzi, Castro-Camilo, Simpson, Lindgren, and Rue (2018) provides an overview of the SPDE approach to spatial modelling with a focus on linking theory to code. Miller, Glennie, and Seaton (2019) summarizes the approach and illustrates its equivalence to penalized smoothing approaches. Lindgren, Bolin, and Rue (2022) provide a recent review of the approach and its applications over the last decade.

For a user of **sdmTMB**, the following are the important elements to understand. First, the SPDE approach links Gaussian random fields (GRFs) that have a Matérn covariance function to Gaussian *Markov* random fields (GMRFs) in such a way that a GMRF can be a good approximation to a GRF (Lindgren *et al*. 2011). This means that GRF models can be computationally approximated as GMRFs. By working with GMRFs, one can take advantage of theory developed to estimate their sparse precision matrix efficiently (Rue and Held 2005; Lindgren *et al*. 2011) and avoid the inversion of large dense matrices. Second, the SPDE involves piece-wise linear basis functions that are defined by a triangulation over the spatial area of interest (Lindgren *et al*. 2011). Commonly, this is referred to as a “mesh”. The properties of this mesh (e.g., its resolution and how far it extends beyond the data) affect the accuracy and computational efficiency of the SPDE approach (Lindgren *et al*. 2011). Third, working with the SPDE approach and the precision matrix of a GMRF (***Q***) introduces an alternative parameter *τ*, which scales the precision matrix, and involves three sparse matrices associated with the mesh (***C, G***_1_, and ***G***_2_) (Lindgren *et al*. 2011),

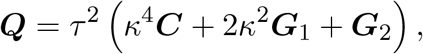

where *κ* is the Matérn decorrelation rate as before. We can calculate the marginal variance of the Matérn random field as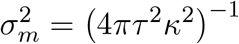.

### 2.5. Adding spatiotemporal random fields

We can extend our spatial model to accommodate spatiotemporal data by adding Gaussian random fields for each time step *t*, ***ϵ***_*t*_

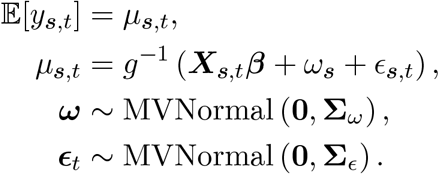

In this case, we assume the spatiotemporal random fields are independent at each time step, but we could alternatively assume they are structured as a random walk or autoregressive process (demonstrated in Section 5.4). The spatiotemporal random fields represent latent variables causing spatial correlation that changes with each time step.

### 2.6. Additional model components

In practice, the above models can become considerably more complex by, for example, letting coefficients vary though time, letting coefficients vary through space (Hastie and Tibshirani 1993) as random fields, or adding random intercepts by grouping factors (Figure 2). All these components are additive in link space. Adopting the notation “main” for main effects, “tvc” for time-varying coefficients, and “svc” for spatially varying coefficients, a more complex model could be written as

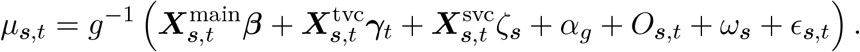

where the ***X*** represent model matrices, ***γ***_*t*_ represents a vector of coefficients that are constrained to vary through time as random walks or AR(1) processes, *ζ*_***s***_ represents spatially varying coefficients following a random field, *α*_*g*_ represents IID random intercepts by group *g* (*α*_*g*_ ∼ Normal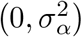), *O*_***s***,*t*_ represents an offset variable (McCullagh and Nelder 1989, p. 206) (e.g., log sampling effort), and *ω*_***s***_ and *ϵ*_***s***,*t*_ represent spatial and spatiotemporal intercept random fields as before (Figure 2). We demonstrate these model components in Sections 4, 5, and 6.

**Figure 2:**
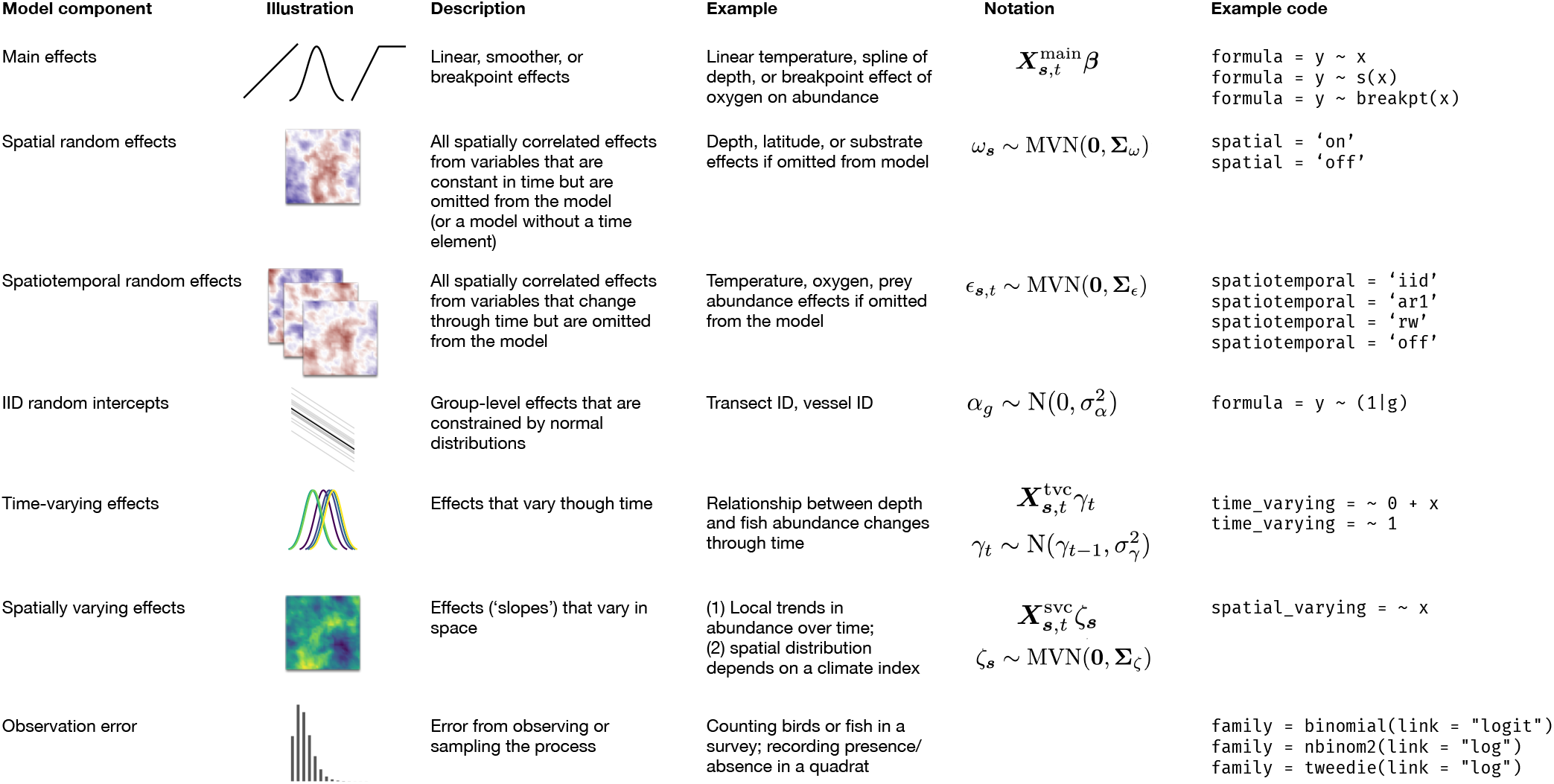
Components of an **sdmTMB** model with illustrations, descriptions, examples, notation, and example code. An **sdmTMB** model can be built from any combination of the process components (first six rows) plus an observation component (last row). The examples are from an ecology context, but the model can be fit to any spatially referenced point data. Notation: We refer to design matrices as ***X***. The indexes ***s***, *t*, and *g* index spatial coordinates, time, and group, respectively. The *σ* and ∑ symbols represent standard deviations and covariance matrices, respectively. All other symbols refer to the described model components (e.g., ***β*** and ***ω*** refer to a vector of main effects and spatial random field deviations, respectively). Note that `s()` denotes a smoother as in **mgcv** (Wood 2017), `breakpt()` denotes a breakpoint “hockey-stick” shape (e.g., Barrowman and Myers 2000), `(1|g)` denotes a random intercept by group `g`, and ∼0 is used in an R formula to omit the intercept.

### 2.7. Delta models

So far, we have described models with one linear predictor and one family (common terminology in R packages for the combination of an observation likelihood and link). Frequently, data are better represented with two-part “delta” or “hurdle” models, which include linear predictors and observation distributions for two processes: zero vs. non-zero values and positive values, respectively (Aitchison 1955). Here, we describe two types of delta models, dropping the space and time subscripts for simplicity.

#### Standard delta models

Using *p* to denote the probability of a non-zero observation and *r* to represent the expected rate for positive data, we can construct two linear predictors (*η*_1_ and *η*_2_) in link space

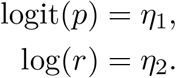

We can relate *p* and *r* to the data via a Bernoulli and positive likelihood distribution (e.g., lognormal or gamma with *ϕ* representing a generic dispersion parameter). We use I(.) to denote an indicator function, which is 1 if *y* > 0 and 0 if *y* = 0.

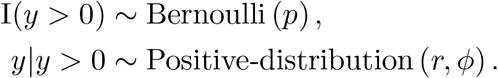

The expectation for a new data point is then the probability of a non-zero event multiplied by the positive rate: *p* · *r*.

#### Poisson-link delta models

An additional delta model is possible that has several advantages over the logit-log delta model (Thorson 2018). The primary advantage is that both linear predictors use a log link so the linear predictors can be added in link space and the partial effect of a coefficient from both linear predictors can be combined. In these models, the linear predictors represent a theoretical group number *n* and a theoretical weight (e.g., mass) per group *w* (Thorson 2018),

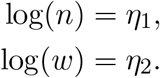

Note that the first linear predictor has a log link, *not* a logit link, and the linear predictor predicts group number *n, not* positive observation probability *p*. These get transformed (Thorson 2018) via

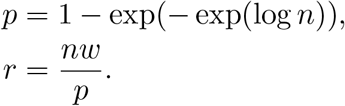

The first part is the complementary-log-log inverse link (McCullagh and Nelder 1989, p. 31) (which has its roots in a Poisson process) but the group number *n* also enters into the expected positive rate *r*. These probabilities *p* and positive rates *r* are then entered into a Bernoulli and positive observation likelihood as before

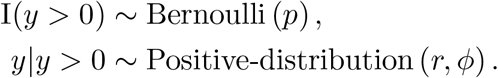

## 3. Software design and user interface

The design goal of **sdmTMB** was to develop a flexible, modular, and intuitive interface to fast maximum likelihood inference (or full Bayesian inference) with the SPDE approach to spatial and spatiotemporal GLMMs with random fields. The package gathers functionality not found combined in other packages that is particularly useful to species distribution modelling, but is applicable beyond ecology to any field that encounters geostatistical data that is continuously referenced in space and (optionally) discretely indexed by time. **sdmTMB** relies on several well-established R packages to construct and fit models (Figure 3).

**Figure 3:**
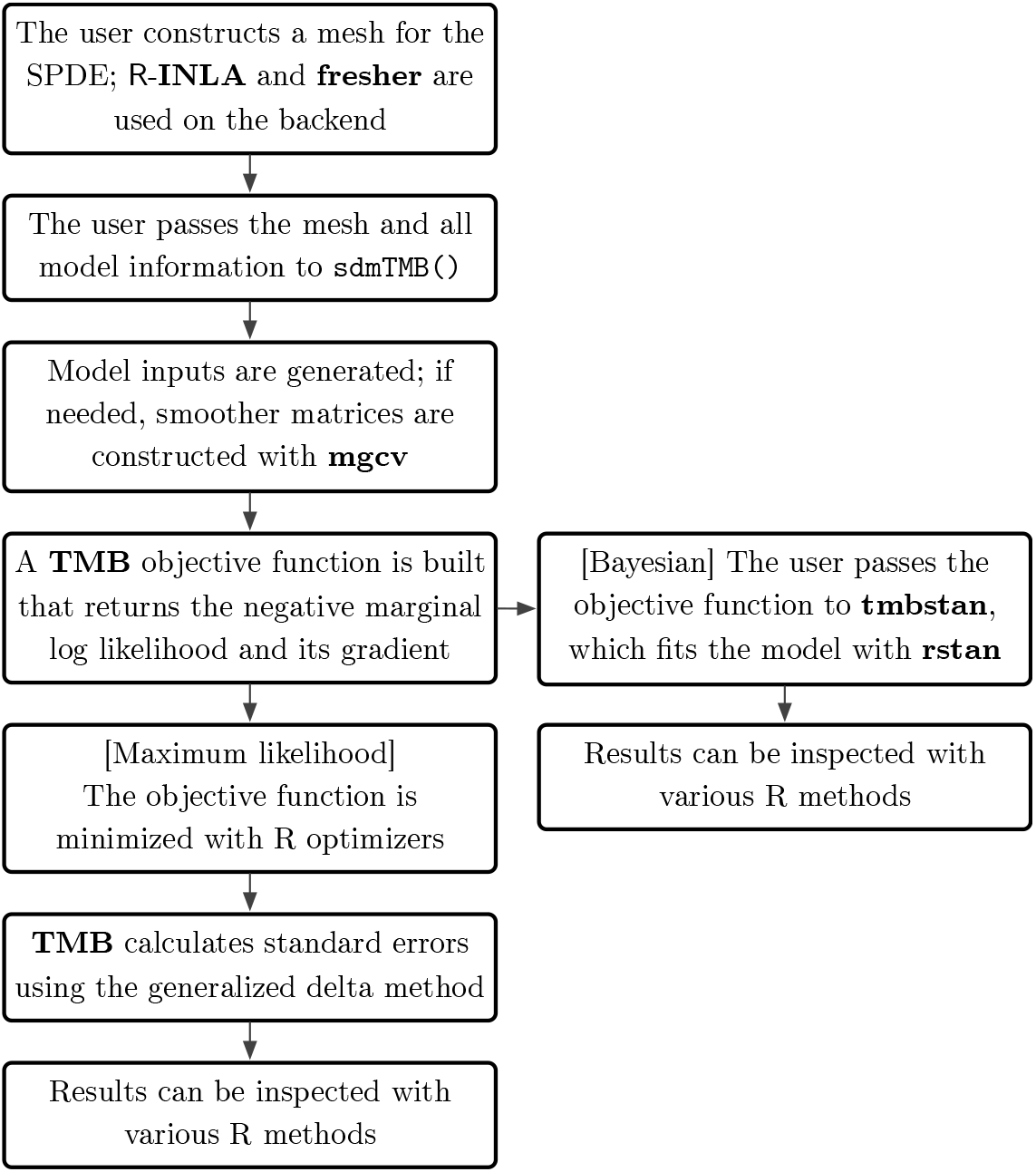
Description of the model fitting procedure in **sdmTMB**.

The **sdmTMB** package is designed to be both modular and familiar to users of widely used R packages (e.g., **glmmTMB**, lme4, **mgcv**). The user starts by constructing a triangulation mesh for the SPDE approach with `make_mesh`() (Figure 3). `make_mesh`() is a wrapper for **fmesher** (Lindgren 2023) triangulation mesh functions and users can also construct any mesh with **fmesher** or R-**INLA** and pass it to `make_mesh`().

Fitting is accomplished with `sdmTMB`(), which has arguments similar to **glmmTMB**’s `glmmTMB`() but with additional arguments for how any spatial and spatiotemporal random fields are structured, what column defines time, any time-varying formulas, and any spatially varying formulas. Observation distributions and links are specified with standard `R` family functions (e.g., `binomial`()) plus several **sdmTMB**-specific families not available in the `R` stats library (e.g., `nbinom2(), delta_lognormal()`, see `?sdmTMB::Families`).

The **sdmTMB** formula syntax (formula argument) follows a standard format that is similar to **glmmTMB, lme4**, and **mgcv**. In addition to standard main effects, the user can include random intercepts (e.g., `+ (1 | group`)), threshold shaped hockey-stick models `breakpt()` (Barrowman and Myers 2000), logistic functions `logistic()`, and penalized smoothers `s()` for generalized additive models, GAMS (Wood 2017). Penalized smoothers use the same `s()` and `t2()` syntax as in mgcv (Wood 2017). Supported functionality includes bivarate smoothers `s(x, y)`, smoothers varying by continuous or categorical variables `s(x1, by = x2)`, cyclical smoothers `s(x, bs = “cc”)`, and smoothers with specified basis dimensions s(x, k = 4) (Wood 2017). Beyond the main formula, `sdmTMB()` accepts one-sided formulas for coefficients that should vary through time (`time_varying`) according to a random walk or AR(1) process (`time_varying_type`) or vary through space as random fields (`spatial_varying`).

Once the user makes a call to `sdmTMB()`, input data structures for a **TMB** model are constructed internally (Figure 3). If needed, data structures required to implement penalized smoothers are constructed with `smooth2random()` from **mgcv** (Wood 2017), similarly to **gamm4** (Wood and Scheipl 2020) and **brms** (Bürkner 2017). `sdmTMB()` formats data, establishes parameter starting values, and constructs an objective function with derivatives based on a compiled C++ template written for TMB. The objective function returns the marginal log likelihood and its gradient, integrating over random effects with the Laplace approximation (Kristensen *et al*. 2016) and efficient using sparse-matrix computation provided by **Matrix** (Bates, Maechler, and Jagan 2024) as an interface to the **Eigen** library in C++. The negative marginal log likelihood is minimized via the non-linear optimization routine `stats::nlminb()` (Gay 1990) in R with additional optimization carried out via a Newton optimization step to find the Hessian using `stats::optimHess()` (Gay 1990; R Core Team 2024). Random effect values are returned at values that maximize the likelihood conditional on the fixed effects at their maximum marginal likelihood (i.e., plug-in or empirical Bayes estimates); however, it is also possible to apply an “epsilon” estimator (Thorson and Kristensen 2016), which corrects for bias arising from the variance and skewness of random effects when calculating an estimator as a non-linear transformation or random effects (see Section 5.7 for an example). Standard errors on all parameters and derived quantities—including those involving random effects—are calculated using the generalized delta method (Kristensen *et al*. 2016; Zheng and Cadigan 2021). After rapid model exploration with maximum likelihood, one can optionally pass an **sdmTMB** model to the R package **rstan** (Carpenter, Gelman, Hoffman, Lee, Goodrich, Betancourt, Brubaker, Guo, Li, and Riddell 2017; Stan Development Team 2022) via **tmbstan** (Monnahan and Kristensen 2018) to sample from the joint posterior distribution for Bayesian inference.

A fitted model summary can be viewed with `print()` or `summary()` and a set of basic “sanity” checks can be run with `sanity()`. `tidy()` returns parameter estimates in standard data frame formats similar to **broom** (Robinson, Hayes, and Couch 2022). Other standard methods are also available such as `fixef()`, `confint()`, and `vcov()`.

Prediction on fitted or new data is accomplished with `predict()` (`?predict.sdmTMB`). In this paragraph, we include relevant `predict()` arguments in parentheses. The predict method is flexible and includes the option to specify a new data frame (`newdata`), whether to return predictions on the link or response scale (`type`), whether to return standard errors (`se_fit`, which can be slow if conditioned on random fields), whether to condition on the random fields (`re_form`), whether to condition on the random intercepts (`re_form_iid`), which delta model linear predictor to use (model), whether to take draws from the joint parameter precision matrix (`nsim`), and whether to use MCMC samples from a **tmbstan** model fit (`mcmc_samples`).

A variety of model evaluation tools are available. A residual`s()` method calculates various types of residuals. The default is a form of randomized quantile (Dunn and Smyth 1996) or probability integral transform (Smith 1985) residuals. New observations can be simulated from a fitted model with `simulate()` or new data can be simulated without a fitted model (“de novo”) with `sdmTMB_simulate()`. `sdmTMB_cv()` facilitates cross validation.

**sdmTMB** models can include penalized likelihoods by assigning priors (penalties) to model parameters through the `sdmTMB()` prior argument (`?sdmTMBpriors`). These priors may be useful in cases where estimation is difficult because of identifiability issues or relatively flat likelihood surfaces, or to impart prior information or achieve regularization. Following other recent SPDE implementations in **TMB** (Breivik, Aanes, Søvik, Aglen, Mehl, and Johnsen 2021; Osgood-Zimmerman and Wakefield 2023), penalized complexity (PC) priors (Simpson, Rue, Riebler, Martins, and Sørbye 2017; Fuglstad, Simpson, Lindgren, and Rue 2019) (`?pc_matern`) can constrain the spatial range and variance parameters. These priors or penalties are available both with maximum likelihood estimation and with MCMC sampling. If a model will be passed to **tmbstan** with priors, a `bayesian` logical argument should be set to `TRUE` to enable Jacobian adjustments for changes of variables (priors applied to parameters that are internally transformed) (Carpenter *et al*. 2017).

### 3.1. Installation

**sdmTMB** can be installed from the Comprehensive R Archive Network (CRAN) at https://CRAN.R-project.org/package=sdmTMB

~~~
*R*> *install.packages*(*“sdmTMB”*)
~~~

Users who wish to automatically install suggested packages as well may wish to use

~~~
*R*> *install.packages*(*“sdmTMB”, dependencies* = *TRUE*)
~~~

As an alternative to the CRAN version, the development version can be installed with

~~~
*R*> *install.packages*(*“pak”*)
*R*> *pak::pkg_install*(*“pbs-assess/sdmTMB”, dependencies* = *TRUE*)
~~~

Additional utilities, which require heavier package dependencies (such as R-**INLA** and **rstan**) and are used by only a subset of users, are maintained in the **sdmTMBextra** package at https://github.com/pbs-assess/sdmTMBextra

Users willing to replace the default R BLAS (Basic Linear Algebra Subprograms) (Blackford, Petitet, Pozo, Remington, Whaley, Demmel, Dongarra, Duff, Hammarling, Henry *et al*. 2002) library with an optimized version (e.g., **openBLAS**; Xianyi, Qian, and Yunquan 2012) can expect up to an order of magnitude increase in model fitting speed for complex models. Suggestions are included in the package README file.

## 4. Example: spatial species distribution modelling

We begin with a simple species distribution model of encounter probability of Pacific Cod (*Gadus macrocephalus*) from a trawl survey conducted in Queen Charlotte Sound, British Columbia, Canada. The purpose of our example is to illustrate the need for spatial random fields. This survey is conducted by Fisheries and Oceans Canada and follows a depth-stratified random sampling design, resulting in a georeferenced dataset. The data frame pcod is available as package data in **sdmTMB**. Relevant columns include latitude, longitude, Universal Transverse Mercator (UTM) coordinates, bottom depth, and encounter (`present = 1`) vs. non-encounter (`present = 0`) of Pacific Cod for a given survey sample.

~~~
*R*> *library*(*sdmTMB*)
*R*> *library*(*dplyr*)
*R*> *library*(*ggplot2*)
*R*> *select*(*pcod, lat, lon, X, Y, depth, present*)
~~~

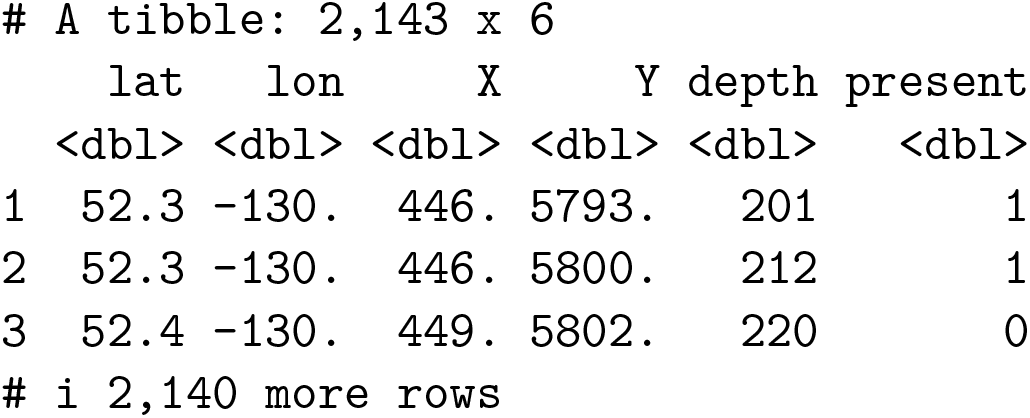

### 4.1. Adding UTM columns

An **sdmTMB** model requires a data frame that contains a response column, columns for any predictors, and columns for spatial coordinates. Usually it makes sense to convert the spatial coordinates to an equidistant projection such as UTMs to ensure that distance remains constant throughout the study region (e.g., using sf:: `st_transform()`, Pebesma 2018). Here we use the helper function `add_utm_columns()` to add UTM coordinates with km units (so our estimated spatial range parameter is not too big or small). By default, the function will guess the UTM zone and create new columns X and Y. Since our example data already has these UTM columns, we can skip running this code.

~~~
*R*> *pcod* <*- add_utm_columns*(*pcod, c*(*“lon”, “lat”*), *units* = *“km”*)
~~~

### 4.2. SPDE mesh creation

We then create a mesh object that contains triangulation and projection matrices needed to apply the SPDE approach using `make_mesh`(). The argument cutoff defines the minimum allowed distance between mesh vertices in the units of X and Y (km). We could create a basic mesh specifying this:

~~~
*R*> *mesh_pcod* <*- make_mesh*(*pcod, xy_cols* = *c*(*“X”, “Y”*), *cutoff* = *8*)
~~~

We can also specify additional arguments, in this case passed to `fmesher::fm_mesh_2d_inla()`: a maximum triangle edge (`max.edge`) length of 10 km and 40 km for the inner and outer mesh, and an `offset` width of 10 km and 40 km for the inner and outer meshes borders. For more irregularly shaped areas, we could have specified a non-convex hull with `convex` and `concave` arguments. The triangle edge length should be several times smaller than the range and the outer boundary should extend at least as far as the range to avoid edge effects. See Krainski *et al*. (2018) chapters 2.6 and 2.7 for guidance on mesh construction.

We can visualize the mesh object with the associated plotting method (Figure 4). Our mesh has 563 (`mesh_pcod2$mesh$n`) vertices. Mesh complexity has a large influence on the speed of fitting these models.

**Figure 4:**
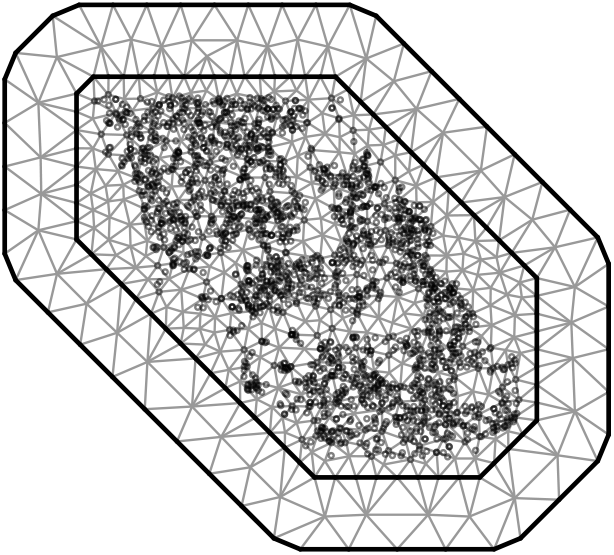
SPDE mesh (lines) combined with the trawl survey observations (points). The locations where lines intersect are referred to as “vertices” or “knots”. Finer meshes will be slower to fit but generally increase the accuracy of the SPDE approximation, to a point. A greater degree of control over the mesh construction can be achieved by using **fmesher** or R-**INLA** directly and supplying the object to `make_mesh`().

~~~
*R*> *mesh_pcod2* <*- make_mesh*(
*+ pcod*,
*+ xy_cols* = *c*(*“X”, “Y”*),
*+ fmesher_func* = *fmesher::fm_mesh_2d_inla*,
*+ cutoff* = *8*,
*+ max.edge* = *c*(*10, 40*),
*+ offset* = *c*(*10, 40*)
*+*)
*R*> *plot*(*mesh_pcod2*)
~~~

### 4.3. Fitting the model

We will fit a logistic regression of encounter probability with and without spatial random fields to illustrate the importance of accounting for spatial correlation. In addition to the spatial random field, we include an intercept and a quadratic effect of depth on the probability of encounter. Our model can be written as

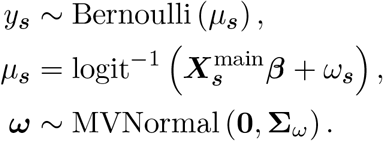

where 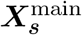 represents a matrix of main effect covariates (intercept and quadratic effects of depth), ***β*** represents a vector of estimated main effect coefficients, and *ω*_***s***_ represents the estimated spatial random field at location ***s***.

We can implement this model with `sdmTMB()`:

~~~
*R*> *fit_bin_rf* <*- sdmTMB*(
*+ present ∼ poly*(*log*(*depth*), *2*),
*+ data* = *pcod*,
*+ mesh* = *mesh_pcod2*,
*+ spatial* = *“on”*,
*+ family* = *binomial*(*link* = *“logit”*)
*+*)
~~~

We can also fit a version that omits the spatial random field by setting `spatial = “off”`. We will use the `update()` method to refit the model while updating any specified arguments:

~~~
*R*> *fit_bin* <*- update*(*fit_bin_rf, spatial* = *“off”*)
~~~

### 4.4. Inspecting the model

We can run some basic checks on our model with the `sanity()` function:

~~~
*R*> *sanity*(*fit_bin_rf*)
#> ✓ Non-linear minimizer suggests successful convergence
#> ✓ Hessian matrix is positive definite
#> ✓ No extreme or very small eigenvalues detected
#> ✓ No gradients with respect to fixed effects are >= 0.001
#> ✓ No fixed-effect standard errors are NA
#> ✓ No fixed-effect standard errors look unreasonably large
#> ✓ No sigma parameters are < 0.01
#> ✓ No sigma parameters are > 100
#> ✓ Range parameter doesn’t look unreasonably large
~~~

This does not flag any issues. `sanity()` is checking that the `nlminb()` optimizer reported successful convergence, that the Hessian matrix is positive definite, that no extreme or small eigenvalues are detected, that no absolute log likelihood gradients with respect to fixed effects are ≥ 0.001, that all fixed effects have reported standard errors that do not look unreasonably large (< 100 by default), that random field marginal standard deviations are not unexpectedly small (< 0.01) or large (> 100), and that the random field Matérn range parameter does not look unreasonably large (≥ 1.5 times the largest distance from a bounding box around the observations).

We can get a summary of our model fit:

~~~
*R*> *summary*(*fit_bin_rf*)
Spatial model fit by ML [‘sdmTMB’]
Formula: present ∼ poly(log(depth), 2)
Mesh: mesh_pcod2 (isotropic covariance)
Data: pcod
Family: binomial(link = ‘logit’)
~~~

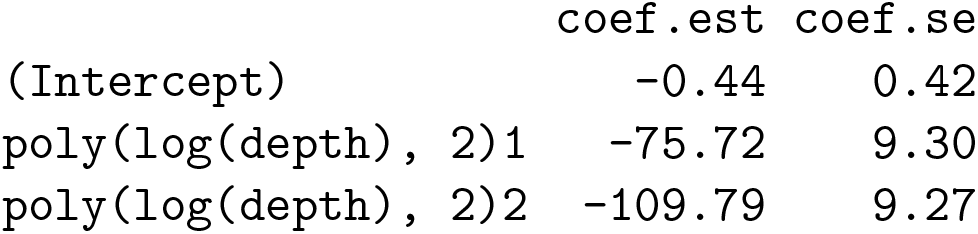

~~~
Matérn range: 41.43
Spatial SD: 1.69
ML criterion at convergence: 1034.744
~~~

See `?tidy.sdmTMB` to extract these values as a data frame.

The output indicates our model was fit by maximum (marginal) likelihood (`ML`). We also see the formula, mesh, fitted data, and family. Next we see any estimated main effects, the Matérn range distance, the spatial random field standard deviation, and the negative log likelihood at convergence.

We can use the `tidy()` function to obtain a data frame with parameter estimates (standard methods such as `fixef()`, `confint()`, and `vcov()` are also available). The standard errors on our fixed effects have increased with the spatial random field:

~~~
*R*> *tidy*(*fit_bin_rf, conf.int* = *TRUE*)
~~~

|  | term | estimate | std.error | conf.low | conf.high |
| --- | --- | --- | --- | --- | --- |
|  | <chr> | <dbl> | <dbl> | <dbl> | <dbl> |
| 1 | (Intercept) | -0.439 | 0.424 | -1.27 | 0.393 |
| 2 | poly(log(depth), 2)1 | -75.7 | 9.30 | -94.0 | -57.5 |
| 3 | poly(log(depth), 2)2 | -110. | 9.27 | -128. | -91.6 |

~~~
*R*> *tidy*(*fit bin, conf.int* = *TRUE*)
~~~

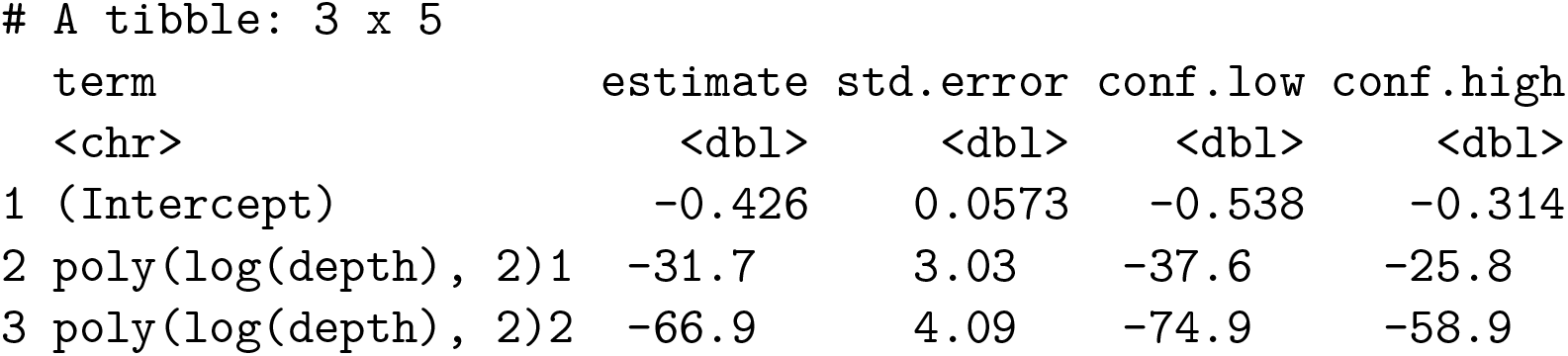

|  | term | estimate | std.error | conf.low | conf.high |
| --- | --- | --- | --- | --- | --- |
|  | <chr> | <dbl> | <dbl> | <dbl> | <dbl> |
| 1 | (Intercept) | -0.426 | 0.0573 | -0.538 | -0.314 |
| 2 | poly(log(depth), 2)1 | -31.7 | 3.03 | -37.6 | -25.8 |
| 3 | poly(log(depth), 2)2 | -66.9 | 4.09 | -74.9 | -58.9 |

By setting `effects = “ran_pars”`, `tidy()` will return random field parameters, where sigma_O is the marginal standard deviation of the spatial random field ***ω*** (“O” for “Omega”).

~~~
*R*> *tidy*(*fit_bin_rf, effects* = *“ran_pars”, conf.int* = *TRUE*)
# A tibble: 2 × 5
 term estimate std.error conf.low conf.high
~~~

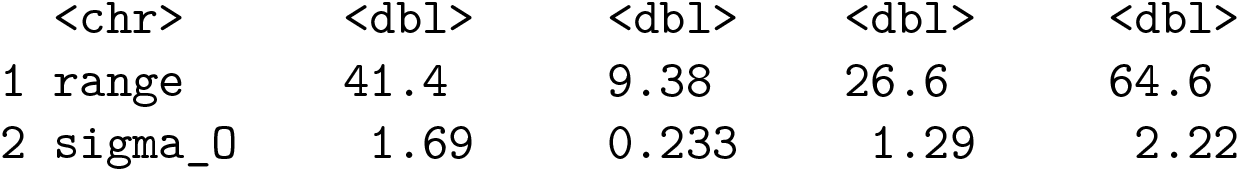

|  | <chr> | <dbl> | <dbl> | <dbl> | <dbl> |
| --- | --- | --- | --- | --- | --- |
| 1 | range | 41.4 | 9.38 | 26.6 | 64.6 |
| 2 | sigma_0 | 1.69 | 0.233 | 1.29 | 2.22 |

### 4.5. Checking the effect of including a random field

We can test for spatial autocorrelation with a visual inspection or a statistical test of the residuals. Here, we demonstrate an approach using an implementation of Moran’s I from the **ape** package (Gittleman and Kot 1990; Paradis and Schliep 2019). We set `type = “mle-mvn”` to denote setting fixed effects at their maximum likelihood estimate (MLE) but taking a single draw from the approximate (multivariate normal) distribution of the random effects (Waagepetersen 2006; Thygesen, Albertsen, Berg, Kristensen, and Nielsen 2017).

~~~
*R*> *inv_dist_matrix* <*- 1 / as.matrix*(*dist*(*pcod[,c*(*“X”, “Y”*), ]))
*R*> *diag*(*inv_dist_matrix*) <*- 0
R*> *set.seed*(*1*)
*R*> *r_bin* <*- residuals*(*fit_bin, type* = *“mle-mvn”*)
*R*> *set.seed*(*1*)
*R*> *r_bin_rf* <*- residuals*(*fit_bin_rf, type* = *“mle-mvn”*)
*R*> *ape::Moran.I*(*r_bin, weight* = *inv_dist_matrix*)*$p.value*
[1] 0
*R*> *ape::Moran.I*(*r_bin_rf, weight* = *inv_dist_matrix*)*$p.value*
[1] 0.8817066
~~~

We see strong evidence for spatial autocorrelation for the model without a random field (p < 0.01) but a lack of evidence for spatial correlation after including a random field suggesting that residual spatial autocorrelation is alleviated by including the random field. The specific p-value is dependent on the seed due to the randomization in the randomized quantile residuals.

We can also see that the marginal Akaike information criterion (AIC) (Akaike 1974) of the model with spatial random fields is lower:

~~~
*R*> *AIC*(*fit_bin_rf, fit_bin*)
~~~

|  | df | AIC |
| --- | --- | --- |
| fit_bin_rf | 5 | 2079.487 |
| fit_bin | 3 | 2392.070 |

Caution is warranted in performing model selection via marginal AIC for models involving penalized smoothing (spatial or otherwise) (e.g. Greven and Kneib 2010; Säfken, Rügamer, Kneib, and Greven 2021). In marginal AIC calculation, the degrees of freedom is based on the number of fixed effects and does not account for the degree of random effect penalization as conditional AIC would. Methods to estimate effective degrees of freedom for similar models were recently demonstrated in Thorson (2024), but are not yet included in **sdmTMB**.

### 4.6. Comparing models with cross validation

As an alternative to AIC, we can conduct model comparison with cross validation. **sdmTMB** includes the helper function `sdmTMB_cv()` to facilitate this. We will do 10-fold cross validation with the folds constructed randomly. We will set the seed each time to ensure the folds are consistent. Using the `fold_ids` argument, we could supply our own folds and conduct spatially blocked cross validation (Roberts, Bahn, Ciuti, Boyce, Elith, Guillera-Arroita, Hauenstein, Lahoz-Monfort, Schröder, Thuiller, Warton, Wintle, Hartig, and Dormann 2017). If we set a parallel plan with the **future** package (Bengtsson 2021), our folds will be fit in parallel.

~~~
*R*> *library*(*future*)
*R*> *plan*(*multisession*)
*R*> *set.seed*(*12928*)
*R*> *cv_bin_rf* <*- sdmTMB_cv*(*present ∼ poly*(*log*(*depth*), *2*),
*+ data* = *pcod, mesh* = *mesh_pcod, spatial* = *“on”*,
*+ family* = *binomial*(), *k_folds* = *10
+*)
*R*> *set.seed*(*12928*)
*R*> *cv_bin* <*- sdmTMB_cv*(*present ∼ poly*(*log*(*depth*), *2*),
*+ data* = *pcod, mesh* = *mesh_pcod, spatial* = *“off”*,
*+ family* = *binomial*(), *k_folds* = *10
+*)
~~~

We can then calculate any performance metric of interest for comparison. A common metric would be the log score or log predictive (likelihood) density (lpd) of the left-out data (Geisser and Eddy 1979; Vehtari, Gelman, and Gabry 2017),

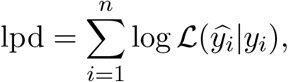

where *n* represents the number of left-out data points, log ℒ is a symbol for log likelihood, *y*_*i*_ represents left-out data point *i*, and *ŷ*_*i*_ represents the prediction for left-out data point *i* with the fixed effects at their MLEs and random effects at their empirical Bayes estimates.

Indeed, the log likelihood predictive density for the left-out data is considerably higher for the model that includes random fields indicating better out-of-sample predictions:

~~~
*R*> *cv_bin_rf$sum_loglik*
[1] -1006.435
*R*> *cv_bin$sum_loglik*
[1] -1195.931
~~~

In practice, we would repeat this procedure several times to ensure the rank order is insensitive to the randomly chosen folds and, if it was, consider averaging across multiple folds or increasing the number of folds.

### 4.7. Making predictions on new data

To visualize our model, we can make predictions with the `predict()` method (`?predict.sdmTMB`) and optionally use the `newdata` argument to predict on a new data frame containing locations and all predictor columns. Here, we will predict on a 2 *×* 2 km grid (qcs_grid) that covers the entire region of interest so we can visualize the predictions spatially. The grid contains spatial covariate columns and all predictors used in the model set at values for which we want to predict. Some covariates might be fixed at a specified value for all predictions, such that we are predicting the expected value for samples conditional on those specified values. In the context of fish or animal surveys, these are sometimes called *detectability* or *catchability* covariates, given the model is predicting the target variable while controlling for the additional influence of these detectability covariates Thorson (2019a). The output of `predict()` is a data frame containing overall estimates in link space (est), estimates from the non-random-field components (`est_non_rf;` here, intercept and depth), and estimates from the individual random field components (here, `omega_s`—the spatial field). We can plot these with `geom_raster()` or `geom_tile()` from the **ggplot2** (Wickham 2016) package (Figure 5).

**Figure 5:**
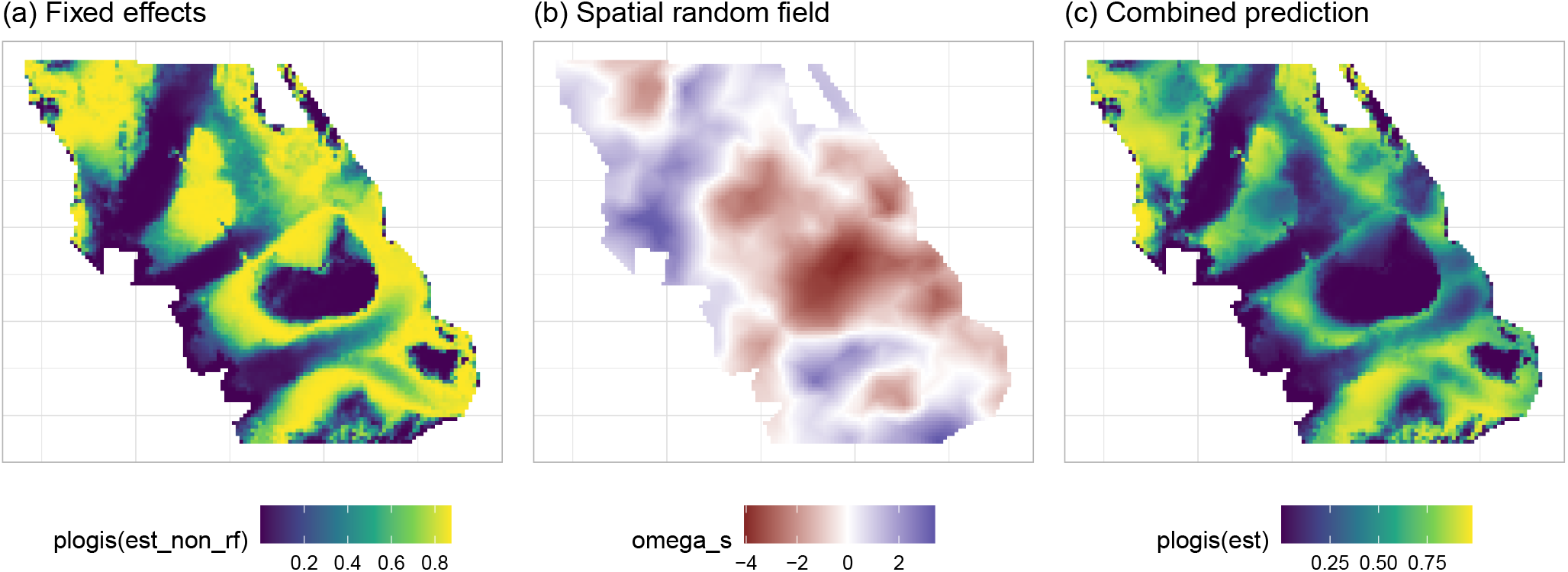
Prediction components from the binomial species distribution model of Pacific Cod. Shown are (a) the quadratic effect of bottom depth, (b) the spatial random field in link (logit) space, and (c) the overall prediction, which here is the combination of panels a and b. The spatial random field represents spatially correlated latent effects not accounted for by the fixed effects. Note the difference between predictions from depth alone (a) and predictions including a spatial random field (c).

~~~
*R*> *p* <*- predict*(*fit_bin_rf, newdata* = *qcs_grid*)
*R*> *select*(*p, X, Y, depth, est, est_non_rf, omega_s*) |>
*+ as_tibble*() |> *head*(*n* = *2*)
~~~

A tibble: 2 x 6
|  | X | Y | depth | est | est_non_rf | omega_s |
| --- | --- | --- | --- | --- | --- | --- |
|  | <dbl> | <dbl> | <dbl> | <dbl> | <dbl> | <dbl> |
| 1 | 456 | 5636 | 347. | -6.52 | -6.23 | -0.291 |
| 2 | 458 | 5636 | 223. | -0.861 | -0.600 | -0.261 |

## 5. Example: spatiotemporal species distribution modeling

As a second example, we will construct a spatiotemporal model of catch rates of Pacific Spiny Dogfish (*Squalus suckleyi*) from a trawl survey off the west coast of Vancouver Island, Canada. This example extends the spatial model described above by including (1) spatiotemporal fields, allowing unique spatially correlated latent effects each year; (2) a time-varying intercept as an AR(1) process, allowing year effects to vary but remain autocorrelated; (3) a smooth effect of depth, allowing catch rates to vary non-linearly with depth; and (4) spatial anisotropy, allowing spatial correlation to be directionally dependent. Since catch rates are positive, continuous, and contain zeros, we begin by specifying the response family as a Tweedie distribution (Tweedie 1984) with a log link. We then compare alternative families, spatiotemporal random field structures, and the exclusion of anisotropy to illustrate the flexibility of **sdmTMB**.

### 5.1. Adding UTM columns and creating a mesh

The dataset includes spatial coordinates, year, dogfish catch weight in kg, area swept in km, and bottom depth:

~~~
*R*> *dat* <*- select*(*dogfish, lon* = *longitude, lat* = *latitude, year*,
*+ catch_weight, area_swept, depth*)
*R*> *dat*
~~~

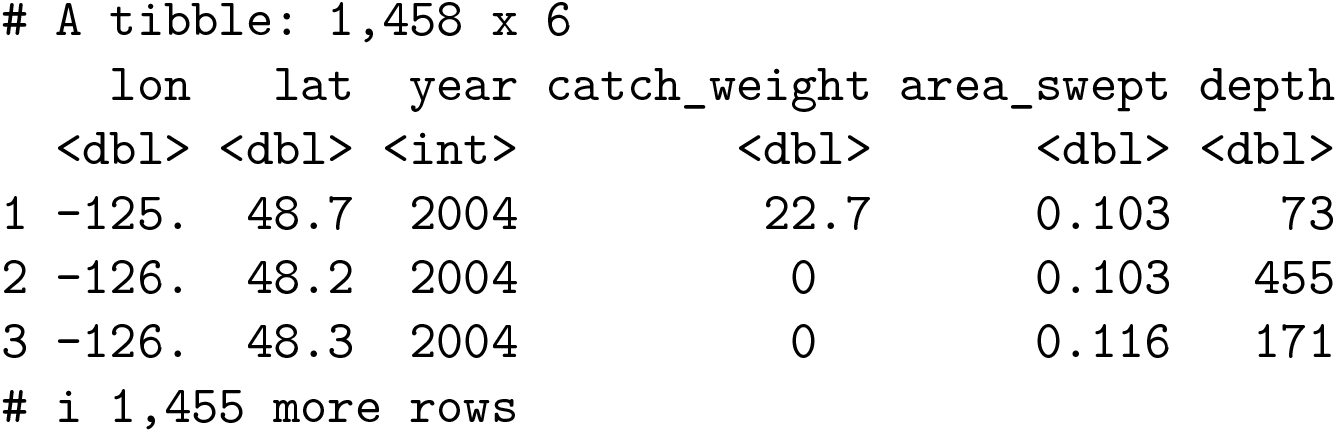

We add UTM zone 9 columns, create a log depth column for convenience, and create a basic mesh:

~~~
*R*> *dat* <*- add_utm_columns*(*dat, c*(*“lon”, “lat”*),
*+ units* = *“km”, utm_crs* = *32609*)
*R*> *dat$log_depth* <*- log*(*dat$depth*)
*R*> *mesh* <*- make_mesh*(*dat, xy_cols* = *c*(*“X”, “Y”*), *n_knots* = *200*)
~~~

### 5.2. Fitting the model

We can then specify our model. We include an offset (McCullagh and Nelder 1989, p. 206) for the effort variable log area swept such that we are effectively modelling density and our predictions will be for an area swept of 1 km^2^.

Our model can be written as

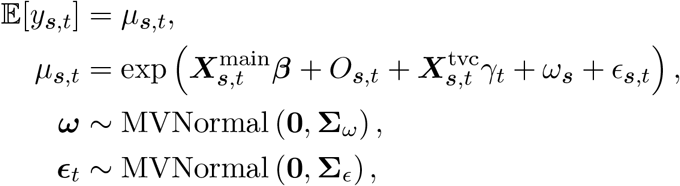

where ***β*** are coefficients associated with the main effects, *O*_***s***,*t*_ represents the offset (here, log area swept), *γ*_*t*_ represents the time-varying coefficients, *ω*_***s***_ is the value from a spatial field (representing constant latent spatial effects) and *ϵ*_***s***,*t*_ is a value from a spatiotemporal field (representing latent spatial effects that vary by year). The temporally varying intercepts *γ*_*t*_ are modeled as a stationary AR(1) process,

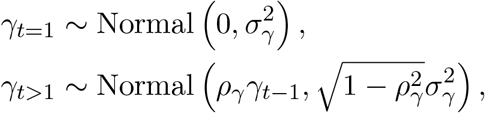

where *ρ*_*γ*_ represents the correlation between intercepts *γ* at time *t*− 1 and *t* and 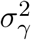 represents the marginal variance of this process. We include the argument extra_time, which represents all time steps to include in the latent process (and all time steps for which we may wish to predict), such that autoregressive processes are applied to equally spaced annual time steps.

We can fit this model as:

~~~
*R*> *fit_tw* <*- sdmTMB*(
*+ catch_weight ∼ s*(*log_depth*),
*+ data* = *dat*,
*+ mesh* = *mesh*,
*+ family* = *tweedie*(),
*+ offset* = *log*(*dat$area_swept*),
*+ time* = *“year”*,
*+ time_varying* = *∼ 1*,
*+ time_varying_type* = *“ar1”*,
*+ spatial* = *“on”*,
*+ spatiotemporal* = *“iid”*,
*+ anisotropy* = *TRUE*,
*+ extra_time* = *seq*(*min*(*dat$year*), *max*(*dat$year*)),
*+ silent* = *FALSE
+*)
~~~

### 5.3. Exploring delta model alternative families

We next explore four alternative families that may better represent the data. Each alternative family uses a delta model formulation as described in Section 2.7.

~~~
*R*> *fit_dg* <*- update*(*fit_tw, family* = *delta_gamma*())
*R*> *fit_dl* <*- update*(*fit_tw, family* = *delta_lognormal*())
*R*> *fit_dpg* <*- update*(*fit_tw, family* = *delta_gamma*(*type* = *“poisson-link”*))
*R*> *fit_dpl* <*- update*(*fit_tw, family* = *delta_lognormal*(*type* = *“poisson-link”*))
~~~

We can then compare the models via AIC:

~~~
*R*> *AIC*(*fit_tw, fit_dg, fit_dl, fit_dpg, fit_dpl*) |>
*+ mutate*(*delta_AIC* = *AIC - min*(*AIC*)) |>
*+ arrange*(*delta_AIC*)
~~~

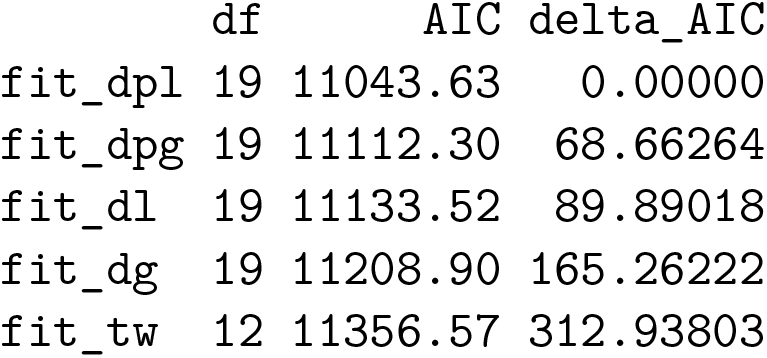

|  | df | AIC | delta_AIC |
| --- | --- | --- | --- |
| fit_dpl | 19 | 11043.63 | 0.00000 |
| fit_dpg | 19 | 11112.30 | 68.66264 |
| fit_dl | 19 | 11133.52 | 89.89018 |
| fit_dg | 19 | 11208.90 | 165.26222 |
| fit_tw | 12 | 11356.57 | 312.93803 |

We find that the Poisson-link delta-lognormal model (Thorson 2018) is favoured by marginal AIC. In an applied situation, we would inspect the distribution of the residuals and consider comparing models with cross validation.

### 5.4. Adding AR(1) random fields and comparing isotropic correlation

We next test two additional model formulations: making the spatial correlation isotropic (the default) instead of anisotropic, and structuring the spatiotemporal random fields as AR(1) to allow spatiotemporal patterns to partially persist from year to year.

The AR(1) fields can be represented as

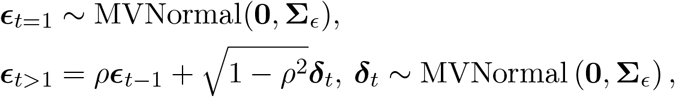

where *ρ* represents the estimated autoregressive parameter allowing the spatial field at time *t* to be correlated with the spatial field at time *t* − 1 with deviations created by ***d***_*t*_, which are themselves independent random fields each year. This is equivalent to a separable model GMRF with precision arising from the Kronecker product of the spatial precision and an AR(1) temporal precision.

~~~
*R*> *fit_dpl_iso* <*- update*(*fit_dpl, anisotropy* = *FALSE*)
*R*> *fit_dpl_ar1* <*- update*(*fit_dpl, spatiotemporal* = *“ar1”*)
*R*> *AIC*(*fit_dpl_ar1, fit_dpl, fit_dpl_iso*)
~~~

|  | df | AIC |
| --- | --- | --- |
| fit_dpl_ar1 | 21 | 11038.88 |
| fit_dpl | 19 | 11043.63 |
| fit_dpl_iso | 17 | 11069.41 |

We find that the anisotropic AR(1) is favoured. It makes sense that anisotropy is important here given the elongated shape of the continental shelf with a rapid transition to deeper water. We can use `plot_anisotropy()` to visually inspect the anisotropy (Figure 6).

**Figure 6:**
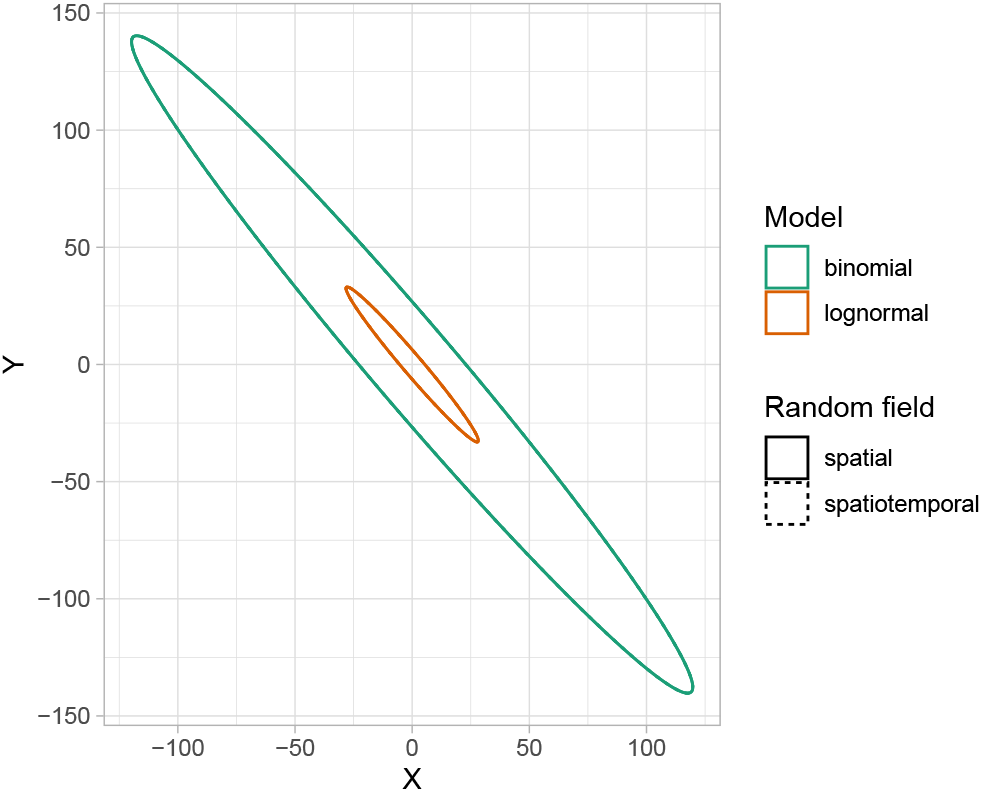
A visualization of anisotropy from the function `plot_anisotropy()`. Ellipses are centered at coordinates of zero in the units that the X-Y coordinates are modeled. The ellipses show the spatial and spatiotemporal range (distance at which correlation is effectively independent) in any direction from the center (zero).

**Figure 7:**
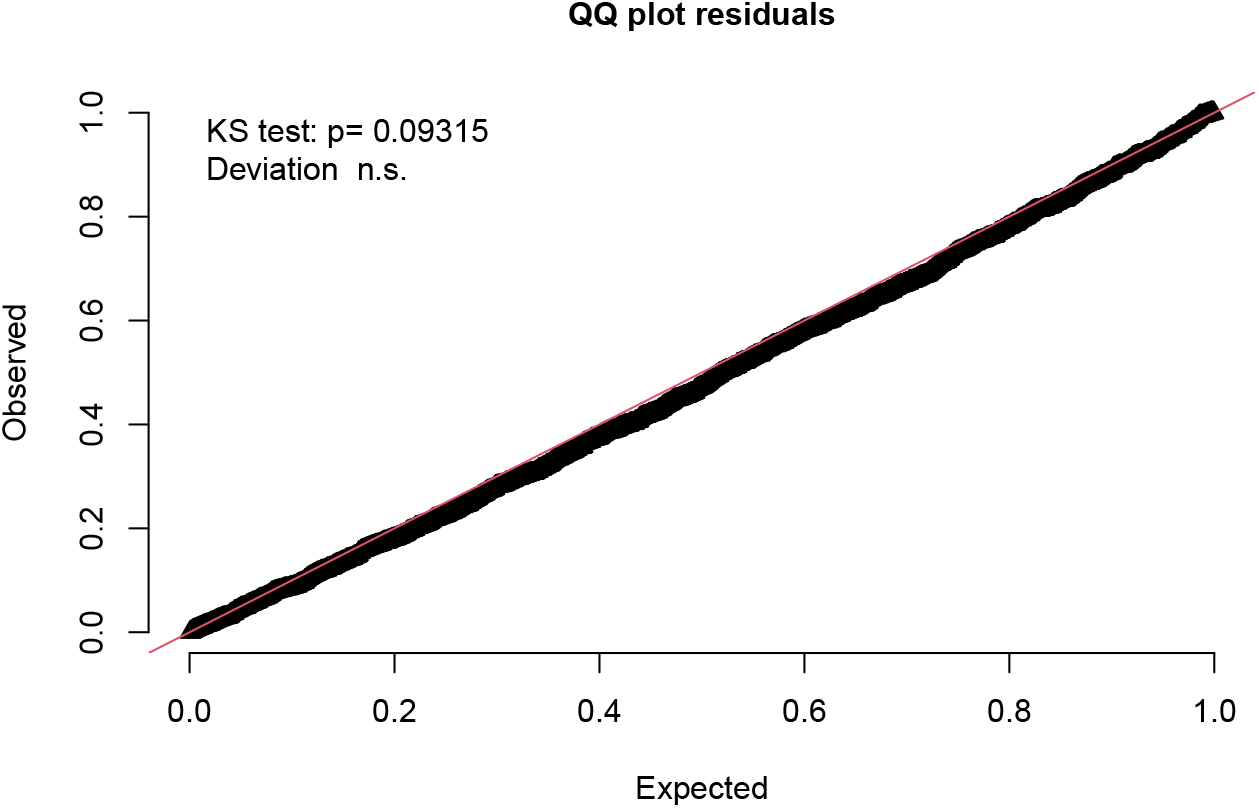
Simulation-based randomized quantile residuals from the **DHARMa** package. The statistical test is a two-sided Kolmogorov-Smirnov test of the null hypothesis that the residuals are drawn from a uniform(0, 1) distribution.

As an example of estimation time for these complex spatiotemporal models, the Tweedie model (`fit_tw`), Poisson-link delta-lognormal model (`fit_dpl`), and Poisson-link delta-lognormal model with autogressive random fields (`fit_dpl_ar1`) took approximately 10, 30, and 90 seconds to fit, respectfully, on an Apple MacBook Pro with an M2 Pro processor and with Apple’s **vecLib** implementation of **BLAS** in R 4.4.0.

~~~
*R*> *plot_anisotropy*(*fit_dpl_ar1*)
~~~

### 5.5. Inspecting the model

We save our chosen model to the object `fit` to simplify subsequent code, run the `sanity()` check (suppressed for brevity), and inspect `summary()`:

~~~
*R*> *fit* <*- fit_dpl_ar1
R*> *sanity*(*fit*)
*R*> *summary*(*fit*)
Spatiotemporal model fit by ML [‘sdmTMB’]
Formula: catch_weight ∼ s(log_depth)
Mesh: mesh (anisotropic covariance)
Time column: year Data: dat
Family: delta_lognormal(link1 = ‘log’, link2 = ‘log’, type = ‘poisson-link’)
Delta/hurdle model 1: -----------------------------------
Family: binomial(link = ‘log’)
~~~

|  | coef.est | coef.se |
| --- | --- | --- |
| (Intercept) | 2.35 | 0.53 |
| slog_depth | 0.20 | 0.67 |

~~~
Smooth terms:
~~~

Smooth terms:
|  | Std. Dev. |
| --- | --- |
| sds(log_depth) | 3.23 |

~~~
Time-varying parameters:
~~~

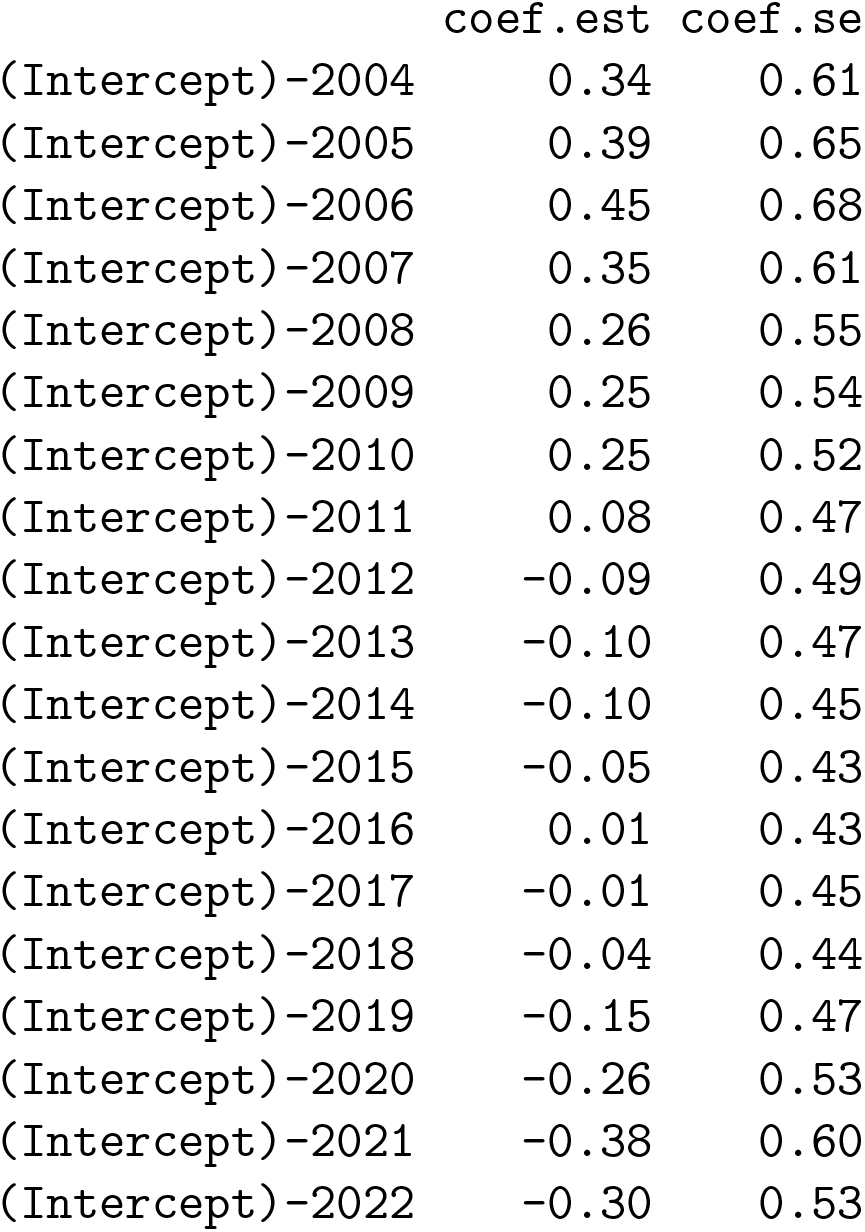

Time-varying parameters:
|  | coef.est | coef.se |
| --- | --- | --- |
| (Intercept)-2004 | 0.34 | 0.61 |
| (Intercept)-2005 | 0.39 | 0.65 |
| (Intercept)-2006 | 0.45 | 0.68 |
| (Intercept)-2007 | 0.35 | 0.61 |
| (Intercept)-2008 | 0.26 | 0.55 |
| (Intercept)-2009 | 0.25 | 0.54 |
| (Intercept)-2010 | 0.25 | 0.52 |
| (Intercept)-2011 | 0.08 | 0.47 |
| (Intercept)-2012 | -0.09 | 0.49 |
| (Intercept)-2013 | -0.10 | 0.47 |
| (Intercept)-2014 | -0.10 | 0.45 |
| (Intercept)-2015 | -0.05 | 0.43 |
| (Intercept)-2016 | 0.01 | 0.43 |
| (Intercept)-2017 | -0.01 | 0.45 |
| (Intercept)-2018 | -0.04 | 0.44 |
| (Intercept)-2019 | -0.15 | 0.47 |
| (Intercept)-2020 | -0.26 | 0.53 |
| (Intercept)-2021 | -0.38 | 0.60 |
| (Intercept)-2022 | -0.30 | 0.53 |

~~~
Spatiotemporal AR1 correlation (rho): 0.74
Matérn anisotropic range (spatial): 17.5 to 183.6 at 130 deg.
Spatial SD: 0.82
Spatiotemporal marginal AR1 SD: 1.34
Delta/hurdle model 2: -----------------------------------
Family: lognormal(link = ‘log’)
~~~

|  | coef.est | coef.se |
| --- | --- | --- |
| (Intercept) | 2.20 | 0.20 |
| slog_depth | 0.11 | 0.24 |

~~~
Smooth terms:
~~~

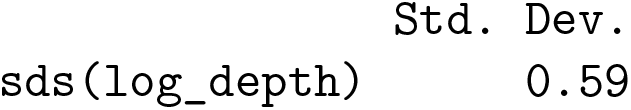

~~~
Time-varying parameters:
~~~

Time-varying parameters:
|  | coef.est | coef.se |
| --- | --- | --- |
| (Intercept)-2004 | 0.43 | 0.27 |
| (Intercept)-2005 | 0.25 | 0.31 |
| (Intercept)-2006 | 0.13 | 0.24 |
| (Intercept)-2007 | 0.03 | 0.30 |
| (Intercept)-2008 | -0.06 | 0.26 |
| (Intercept)-2009 | 0.20 | 0.30 |
| (Intercept)-2010 | 0.50 | 0.26 |
| (Intercept)-2011 | 0.30 | 0.30 |
| (Intercept)-2012 | 0.18 | 0.23 |
| (Intercept)-2013 | 0.10 | 0.29 |
| (Intercept)-2014 | 0.04 | 0.23 |
| (Intercept)-2015 | -0.10 | 0.29 |
| (Intercept)-2016 | -0.26 | 0.23 |
| (Intercept)-2017 | -0.24 | 0.30 |
| (Intercept)-2018 | -0.27 | 0.23 |
| (Intercept)-2019 | -0.26 | 0.35 |
| (Intercept)-2020 | -0.30 | 0.35 |
| (Intercept)-2021 | -0.41 | 0.24 |
| (Intercept)-2022 | -0.20 | 0.25 |

~~~
Dispersion parameter: 1.10
Spatiotemporal AR1 correlation (rho): 0.22
Matérn anisotropic range (spatial): 4.1 to 43.3 at 130 deg.
Spatial SD: 0.41
Spatiotemporal marginal AR1 SD: 0.82
ML criterion at convergence: 5498.439
See ?tidy.sdmTMB to extract these values as a data frame.
See ?plot_anisotropy to plot the anisotropic range.
~~~

The output is more complex compared to our binomial spatial model. We now have two model components, which are shown one after the other. Starting with the binomial component, we have output from the smoother, which includes a linear parameter (`slog_depth`) and the standard deviation on the smoother weights (`sds(log_depth`)). The smoother summary follows the format used in the **brms** package (Bürkner 2017). Next, we have the time-varying intercepts and information on our anisotropic spatial correlation. We then have the second model component (lognormal) with a similar summary structure but with the addition of a dispersion parameter, the AR(1) correlation of the spatiotemporal random fields, and a spatiotemporal random field marginal standard deviation.

We can check simulation-based randomized quantile residuals from our chosen model via the **DHARMa** package (Hartig 2021). To do that, we simulate from our model with the simulate.`sdmTMB()` method and pass those simulations to a helper function `dharma_residuals()`.

~~~
*R*> *set.seed*(*42*)
*R*> *s* <*- simulate*(*fit, nsim* = *500, type* = *“mle-mvn”*)
*R*> *dharma_residuals*(*s, fit, test_uniformity* = *TRUE*)
~~~

The quantile-quantile plot suggests that under the model assumptions, the (transformed) residuals are reasonably consistent with an independent uniform(0, 1) distribution.

### 5.6. Visualizing model predictions

Similarly to the first example, we can visualize model predictions on a grid covering the area of interest. Because this is a spatiotemporal model, we first need to replicate our grid for each year we will predict on. Since this is a common operation, we include the function `replicate_df()` to replicate a data frame. We then ensure our data frame contains all the predictors used in the model (here `log_depth`).

~~~
*R*> *grid* <*- replicate_df*(*wcvi_grid, “year”, time_values* = *unique*(*dat$year*))
*R*> *grid$log_depth* <*- log*(*grid$depth*)
*R*> *head*(*grid, n* = *2*)
~~~

|  | X | Y | depth | year | log_depth |
| --- | --- | --- | --- | --- | --- |
| 1 | 733.9852 | 5346.334 | 447 | 2004 | 6.102559 |
| 2 | 735.9852 | 5346.334 | 442 | 2004 | 6.091310 |

~~~
*R*> *pred* <*- predict*(*fit, newdata* = *grid, type* = *“response”*)
*R*> *names*(*pred*)
~~~

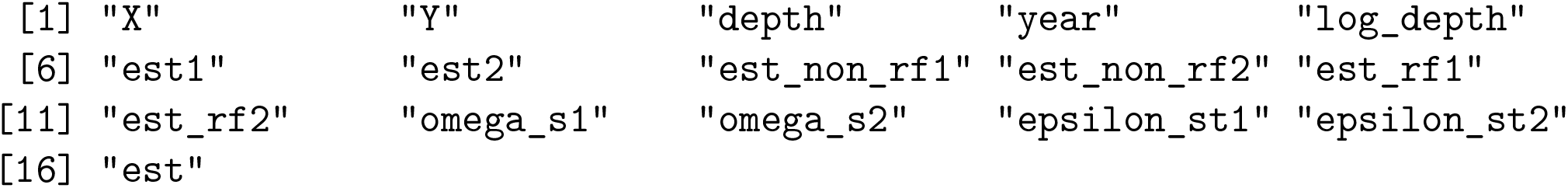

Our prediction data frame is similar to the binomial spatial model, but includes columns for the two delta model linear predictors (labelled with suffixes 1 and 2) and adds an `epsilon_st` column for spatiotemporal random effects. We can easily generate plots from this data frame using **ggplot2** code with `geom_raster()` similarly to our spatial example with Pacific Cod (Figure 8). We suppress that code for brevity.

**Figure 8:**
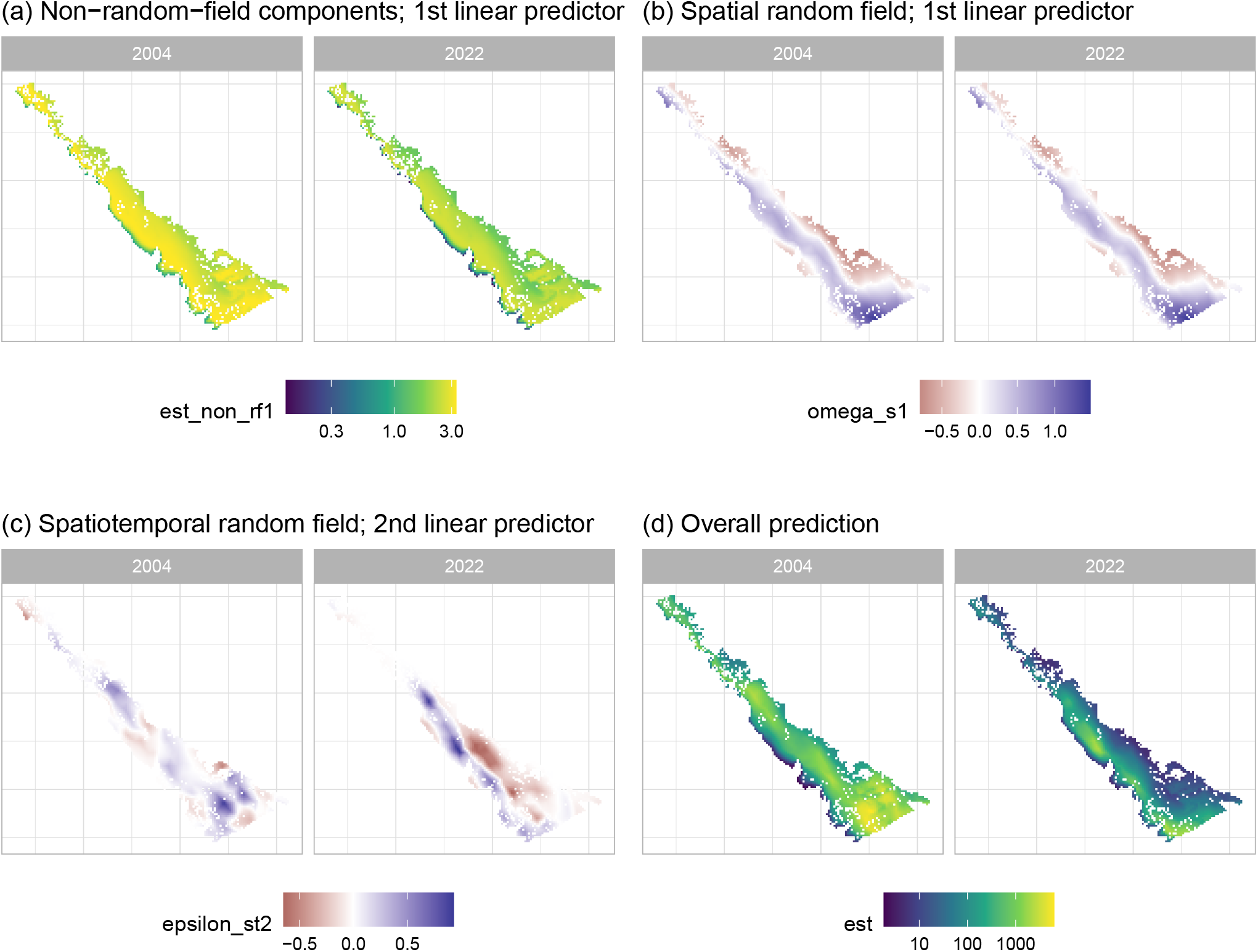
Example prediction elements from the spatiotemporal model of Pacific Dogfish biomass density. Throughout, two example years are shown. (a) `est_non_rf1` refers to the prediction from all non-random-field elements (here, a smoother for bottom depth and the time-varying year effect) from the first linear predictor, (b) `omega_s1` refers to the spatial random field from the first linear predictor, (c) `epsilon_st2` refers to spatiotemporal random fields from the second linear predictor, and (d) `est` refers to the overall prediction estimate combining all effects. The spatial random field is constant through time (i.e., the two panels in b are identical) and represents static biotic or abiotic features not included as covariates (e.g., habitat). The spatiotemporal random fields are different each time step and here are constrained to follow an AR(1) process. They represent temporal variability in the spatial patterning of Pacific Spiny Dogfish (e.g., resulting from movement or local changes in population density).

We can visualize the conditional effect of the bottom depth smoother by predicting across a sequence of depths and holding other variables at reference values (Figure 9). Here, we pick the last year, specify to include both delta model components (`model = NA`), omit the random fields (`re_form = NA`), and return standard errors (`se_fit = TRUE`). Alternatively, we could produce a similar plot using **ggeffects** (Lüdecke 2018) with ggeffects::ggpredict().

**Figure 9:**
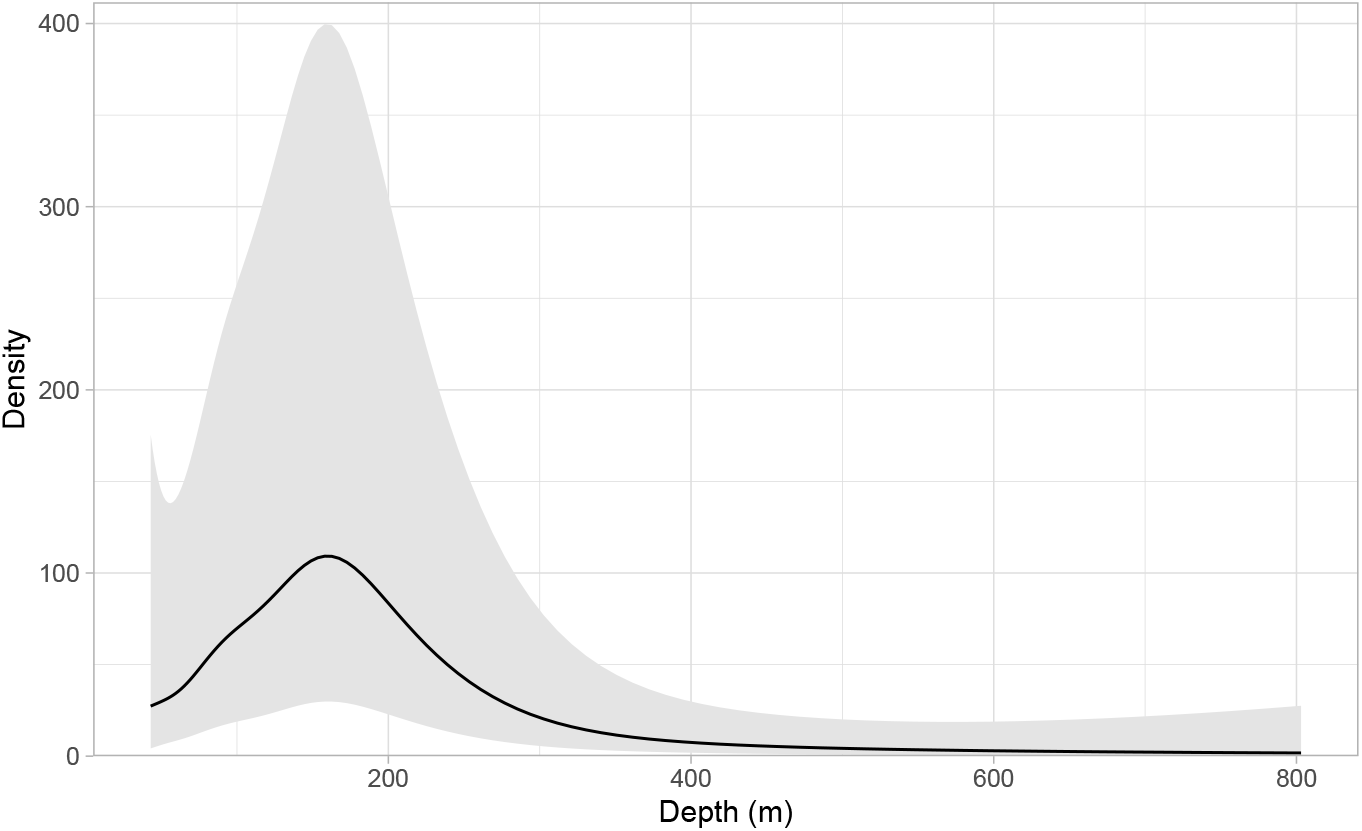
The conditional effect of ocean bottom depth on Pacific Spiny Dogfish population density. The line and shaded ribbon represent the mean and 95% confidence interval, respectively. Other fixed effects are held at constant values and the random fields are set to their expected value (zero).

~~~
*R*> *nd* <*- data.frame*(
*+ log_depth* = *seq*(*min*(*dat$log_depth*), *max*(*dat$log_depth*), *length.out* = *100*),
*+ year* = *max*(*dat$year*)
*+*)
*R*> *pred_depth* <*- predict*(
*+ fit, newdata* = *nd*,
*+ model* = *NA, re_form* = *NA, se_fit* = *TRUE
+*)
~~~

### 5.7. Calculating an area-weighted index

We can generate an area-weighted population index (e.g., a relative or absolute index of abundance or biomass) that is independent of sampling locations by predicting from the model on a grid covering the area of interest and summing the predicted biomass with the `get_index()` function (Figure 10). We supply the grid cell area (4 km^2^) to the area argument and specify `bias_correct = TRUE` to enable a bias correction needed due to the non-linear transformation of the random effects (Thorson and Kristensen 2016).

**Figure 10:**
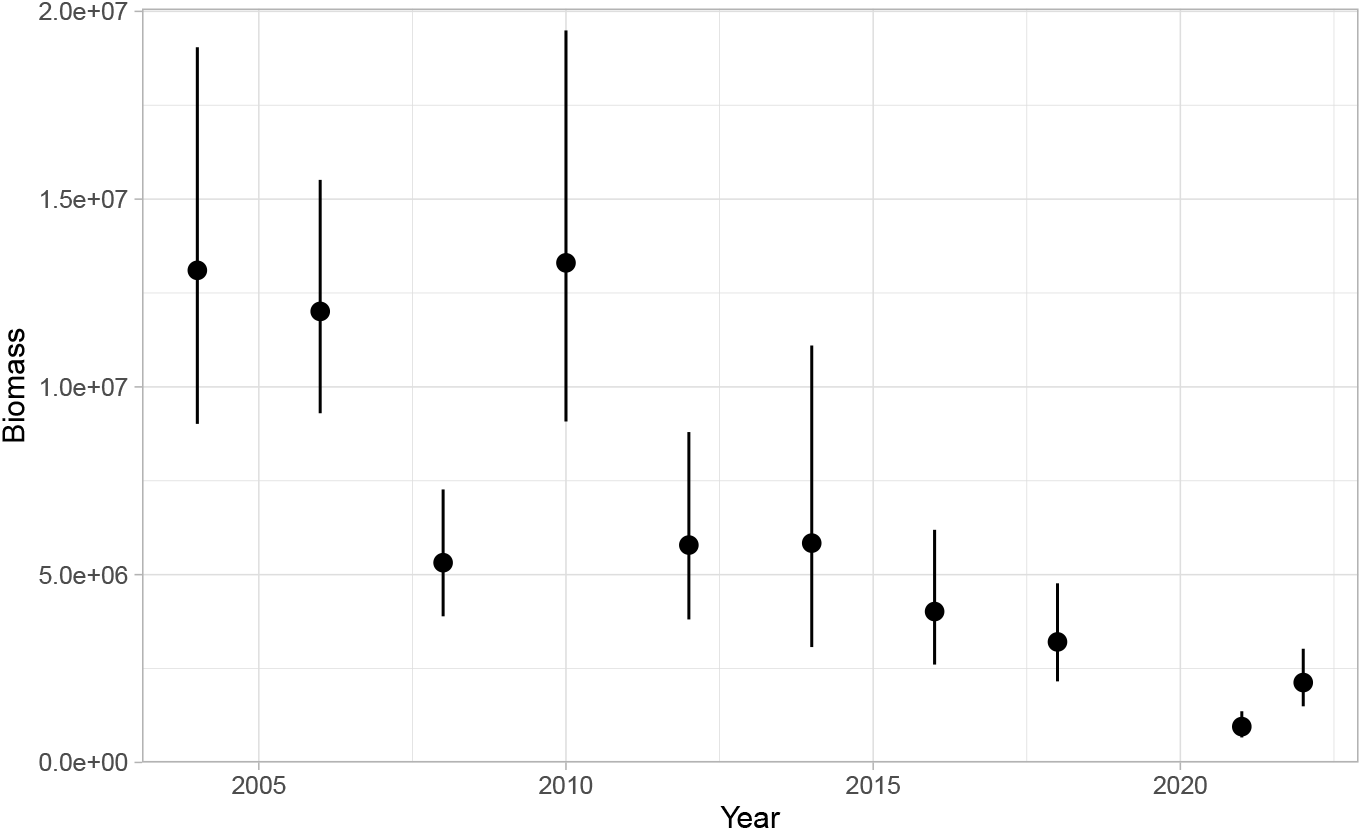
Area-weighted index of relative biomass over time for Pacific Spiny Dogfish. Dots and line segments represent means and 95% confidence intervals.

~~~
*R*> *grid$area* <*- 4 # 2 km x 2 km
R*> *pred2* <*- predict*(*fit, newdata* = *grid, return_tmb_object* = *TRUE*)
*R*> *ind* <*- get_index*(*pred2, bias_correct* = *TRUE, area* = *grid$area*)
~~~

## 6. Example: spatially varying coefficients

In our final example, we demonstrate a model with spatially varying coefficient (SVC) effects and illustrate combining uncertainty from parameters by working with draws from the joint parameter precision matrix. SVC models are a class of models in which coefficients are allowed to vary spatially constrained by some smooth function (Hastie and Tibshirani 1993; Thorson, Barnes, Friedman, Morano, and Siple 2023).

Snowy Owls (*Bubo scandiacus*) breed on the arctic tundra and are irruptive migrants, meaning that they appear across the mid-latitudes of North America in much greater numbers in some winters than others. The reasons for this interannual variation in the number of individuals migrating south are not well understood but seem to be related to high abundances of food during the breeding season and therefore sharp increases in breeding ground population densities (Robillard, Therrien, Gauthier, Clark, and Bêty 2016). The North Atlantic Oscillation Index (NAO) has been linked to productivity of both owls and their prey in Europe (Millon, Petty, Little, Gimenez, Cornulier, and Lambin 2014). Because both productivity and the choice of wintering location could be influenced by climate, we model an SVC effect of annual mean NAO index on early winter abundance across the southern boundary of their winter distribution. Annual mean NAO captures the preceding winter’s conditions, combined with breeding season and early winter climate.

To do this, we use counts of Snowy Owls observed by the annual Christmas Bird Counts (National Audubon Society 2021) from all locations where they have been recorded and for which there were at least three counts conducted between 1979 and 2020. Our data are read from supplementary data and also contain columns for spatial coordinates (in an Albers projection for North America and divided by 100000 to give units of 100 km), year, year as a factor, the annual NAO value, and owl count:

~~~
*R*> *select*(*snow, X, Y, year, year_f, nao, count*) |> *head*()
~~~

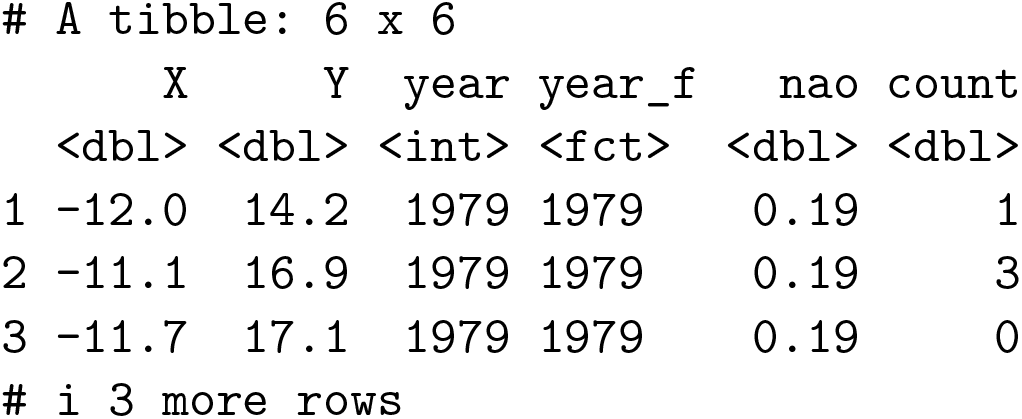

### 6.1. Fitting the model

We will fit counts using a negative binomial (NB2; Hilbe 2011) distribution, random intercepts for year, spatial and spatiotemporal random fields, and an SVC associated with the NAO. Centering and scaling variables (e.g., by their mean and standard deviation) can be helpful to reduce the correlation between the SVC and other random fields that are included in the model when estimating SVCs. Here, NAO is an index with a mean near zero and an SD not too far from 1, so we will leave it as is. However, we also include NAO as a main effect since the SVC random field is drawn from a multivariate normal distribution with mean zero.

We can write the model as

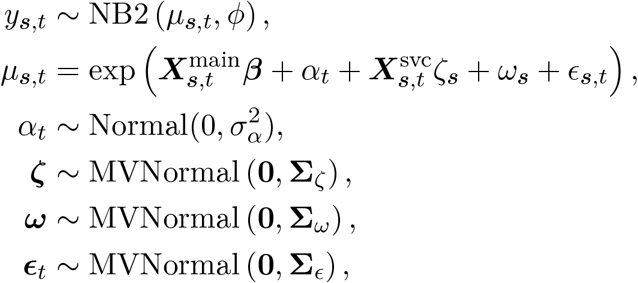

where three types of random fields are now included: spatially varying NAO coefficients (*ζ*_***s***_), a spatial intercept (*ω*_***s***_), and spatiotemporal variation (*ϵ*_***s***,*t*_). The NB2 distribution is specified with a mean *µ*_***s***,*t*_ and size parameter *ϕ*. The observation variance scales quadratically with the mean: Var[*y*] = *µ* + *µ*^2^*/ϕ* (Hilbe 2011). The *α*_*t*_ represent IID random intercepts by year. We can then fit this model:

~~~
*R*> *mesh_snow* <*- make_mesh*(*snow, xy_cols* = *c*(*“X”, “Y”*), *cutoff* = *1.5*)
*R*> *fit_owl* <*- sdmTMB*(
*+  count ∼ 1 + nao +* (*1* | *year_f*),
*+  spatial_varying* = *∼ 0 + nao*,
*+  time* = *“year”*,
*+  data* = *snow*,
*+  mesh* = *mesh_snow*,
*+  family* = *nbinom2*(*link* = *“log”*),
*+  spatial* = *“on”*,
*+  spatiotemporal* = *“iid”*,
*+  silent* = *FALSE
+*)
~~~

### 6.2. Inspecting the model

`summary()` prints standard model information:

~~~
*R*> *summary*(*fit_owl*)
Spatiotemporal model fit by ML [‘sdmTMB’]
Formula: count ∼ 1 + nao + (1 | year_f)
Mesh: mesh_snow (isotropic covariance)
Time column: year
Data: snow
Family: nbinom2(link = ‘log’)
~~~

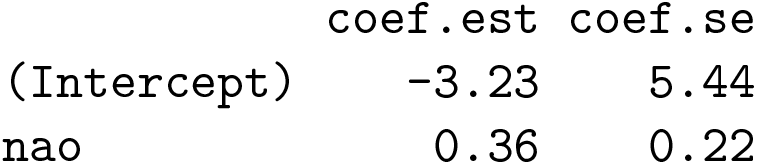

~~~
Random intercepts:
Std. Dev.
year_f 0.3
Dispersion parameter: 0.46
Matérn range: 25.21
Spatial SD: 6.97
Spatially varying coefficient SD (nao): 0.17
Spatiotemporal IID SD: 0.79
ML criterion at convergence: 17512.366
See ?tidy.sdmTMB to extract these values as a data frame.
~~~

In addition to the output seen for other models, we now have a section for random intercepts and a standard deviation for the SVC random field. Given our model specification, all random fields are sharing a single Matérn range. We can also check the confidence intervals (CIs) on the main effect of NAO and see that they overlap zero.

~~~
*R*> *tidy*(*fit_owl, conf.int* = *TRUE*)
~~~

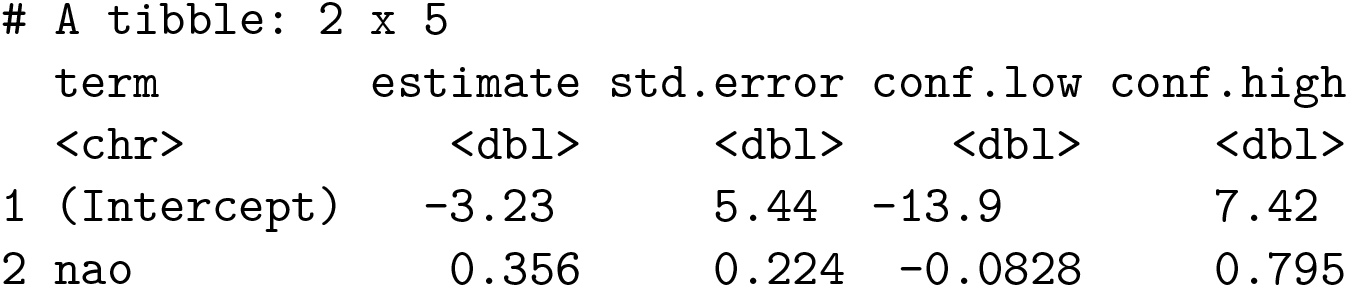

However, given that this is only part of the effect of NAO that we include in our model, we next assess if this variable has a meaningful effect in some locations, even if not overall.

### 6.3. Extracting the spatially varying effects with uncertainty

The spatially varying effect at any point in space is the combination of the main effect and SVC random effect for nao. Mean estimates of the SVC random effect are available in the output of `predict.sdmTMB()` in a column starting with `zeta_s` (in this case, `zeta_s_nao`). However, we might wish to combine the fixed and random components of a spatially varying effect and assess the uncertainty of these combined predictions. We illustrate a way of accomplishing this by simulating from the fixed and random effects while assuming that parametric uncertainty is well approximated using a multivariate normal distribution and the joint precision matrix. We do this by specifying a non-null number of simulation draws to `nsim` in `predict.sdmTMB()`. By default, nsim > 0 will return a matrix of draws for the overall prediction. Here, we instead specify that we want to return draws for the `zeta_s` (*ζ*_***s***_) random field, which is the SVC random field (`sims_var = “zeta_s”`). This returns a matrix where each row matches a row of newdata and each column is a simulation draw. We then use the function spread_sim`s()` to draw 200 simulations for the parameters themselves. Because the simulations are stored in different dimensions, the random field draws must be transposed `t()` before combining the vector of main effect draws (`sims$nao`) with the random field values `zeta_s`. Next, we can calculate the median, and upper and lower quantiles for each column of data, which correspond to the rows in the data provided. For this example, we use thresholds of 0.025 and 0.975 representing a 95% CI.

~~~
*R*> *set.seed*(*42*)
*R*> *zeta_s* <*- predict*(*fit_owl, newdata* = *snow, nsim* = *300, sims_var* = *“zeta_s”*)
*R*> *dim*(*zeta_s*)
[1] 30392 300
*R*> *sims* <*- spread_sims*(*fit_owl, nsim* = *300*)
*R*> *dim*(*sims*)
[1] 300 9
*R*> *combined* <*- sims$nao + t*(*zeta_s*)
*R*> *snow$nao_effect* <*- exp*(*apply*(*combined, 2, median*))
*R*> *snow$nao_effect_lwr* <*- exp*(*apply*(*combined, 2, quantile, probs* = *0.025*))
*R*> *snow$nao_effect_upr* <*- exp*(*apply*(*combined, 2, quantile, probs* = *0.975*))
~~~

We can make a basic plot using the following code. A more elaborate version including separate panels for each of the CIs is shown in Figure 11.

**Figure 11:**
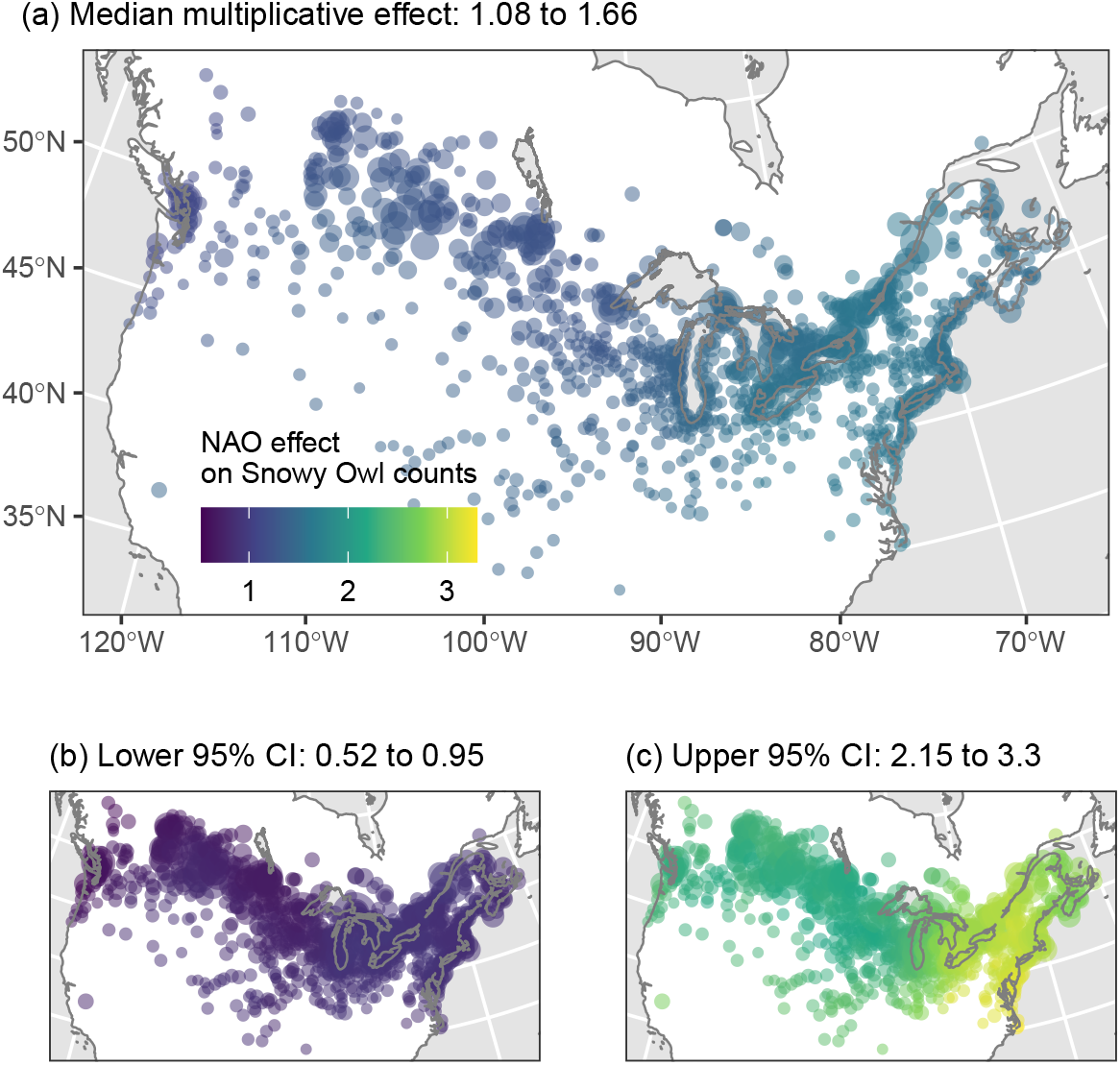
Spatially varying effect of mean annual NAO (North Atlantic Oscillation) on counts of Snowy Owls observed on annual Christmas Bird Counts from 1979–2020 in Canada and the US. The effect is multiplicative on owl count per NAO unit. In the west, the lower bound of values overlaps 1 implying no effect, whereas in the southeast the effect becomes positive. Point size is scaled to the mean counts in each location.

~~~
*R*> *ggplot*(*snow, aes*(*X, Y*)) *+ geom_point*(*aes*(*colour* = *nao_effect*))
~~~

Overall, we find a weak average positive effect of annual mean NAO on overall counts with a southeast to northwest gradient in the intensity of the effect (Figure 11). At some locations, the lower CI on the exponentiated effect is above 1. This result is consistent with owls closest to the Atlantic coast and those migrating the furthest south being the most affected by the NAO.

## 7. Package comparisons

There are many R packages capable of fitting geostatistical spatial or spatiotemporal models (e.g., Heaton *et al*. 2019). **sdmTMB, VAST, tinyVAST, R-INLA/inlabru**, and **spaMM** (Rousset and Ferdy 2014) are the most closely related, as they all provide a user interface to SPDE-based GMRF models. In our software comparison (Table 1), we also include **mgcv** as it can be adapted to use the SPDE (Miller *et al*. 2019) and **spBayes** (Finley, Banerjee, and Carlin 2007; Finley, Banerjee, and E. Gelfand 2015) since it is a prominent package that can fit related predictive-process models without the SPDE. **sdmTMB, VAST**, and **mgcv** can estimate anisotropic covariance whereas R-INLA/**inlabru** and **spBayes** are currently limited to isotropic covariance. To our knowledge, **VAST** and **tinyVAST** are the only packages to implement spatial (Thorson, Scheuerell, Shelton, See, Skaug, and Kristensen 2015a) and spatial dynamic factor analysis (Thorson, Ianelli, Larsen, Ries, Scheuerell, Szuwalski, and Zipkin 2016) and spatial empirical orthogonal function (EOF) regression (Thorson, Cheng, Hermann, Ianelli, Litzow, O’Leary, and Thompson 2020). Of these packages, only **sdmTMB** and **inlabru** can currently fit threshold (e.g., hockey-stick) covariate relationships. To our understanding, **spaMM** is limited to a spatial random field (i.e., does not fit spatiotemporal fields) and **spBayes** implements spatiotemporal fields, but only as a random walk. There is considerable variability in the available observation likelihoods across packages (Table 1).

We ran a simple speed comparison between **sdmTMB**, R-INLA/**inlabru, spaMM**, and **mgcv** for fitting an SPDE spatial random field model to 1,000, 10,000, or 100,000 data points with Gaussian error across a range of mesh resolutions (Figure 12, Appendix A). Our test was restricted to one core and default R algebra libraries using R 4.4.0 and Matrix version 1.7.0. With up to 10,000 rows of data, **sdmTMB** was fastest at approximately a three to 13-fold speed increase over R-INLA/**inlabru** (Figure 12a–b). At larger sample sizes, **inlabru** was more affected by mesh resolution than **sdmTMB** (Figure 12c). **mgcv** was most affected by mesh resolution (Figure 12)—timing was on par or faster than the other packages at low (≈ 200) mesh resolutions but much slower as mesh complexity grew into commonly used ranges. The **mgcv** model was fit with the `bam()` function instead of the standard gam(). The `bam()` function here uses numerical methods optimized for large datasets, including a discretization of covariate values (Wood, Goude, and Shaw 2015; Wood, Li, Shaddick, and Augustin 2017; Li and Wood 2020). The standard `gam()` function was approximately 60 times slower than `bam()` for the largest dataset with the most complex mesh (not shown). **spaMM** scaled well to large datasets with high mesh resolutions (Figure 12c), but to our knowledge can only fit spatial GMRFs. At low mesh resolutions, a large portion of the differences in timing are likely a result of initial model processing and not estimation itself. Speed increases can allow for more rapid and thorough model exploration and experimentation with this class of computationally intensive models. For users ultimately interested in Bayesian inference, the approximate Bayesian inference offered by R-INLA/**inlabru** is likely to be faster than passing the same model from **sdmTMB**/VAST to **tmbstan** for full MCMC-based Bayesian inference.

**Figure 12:**
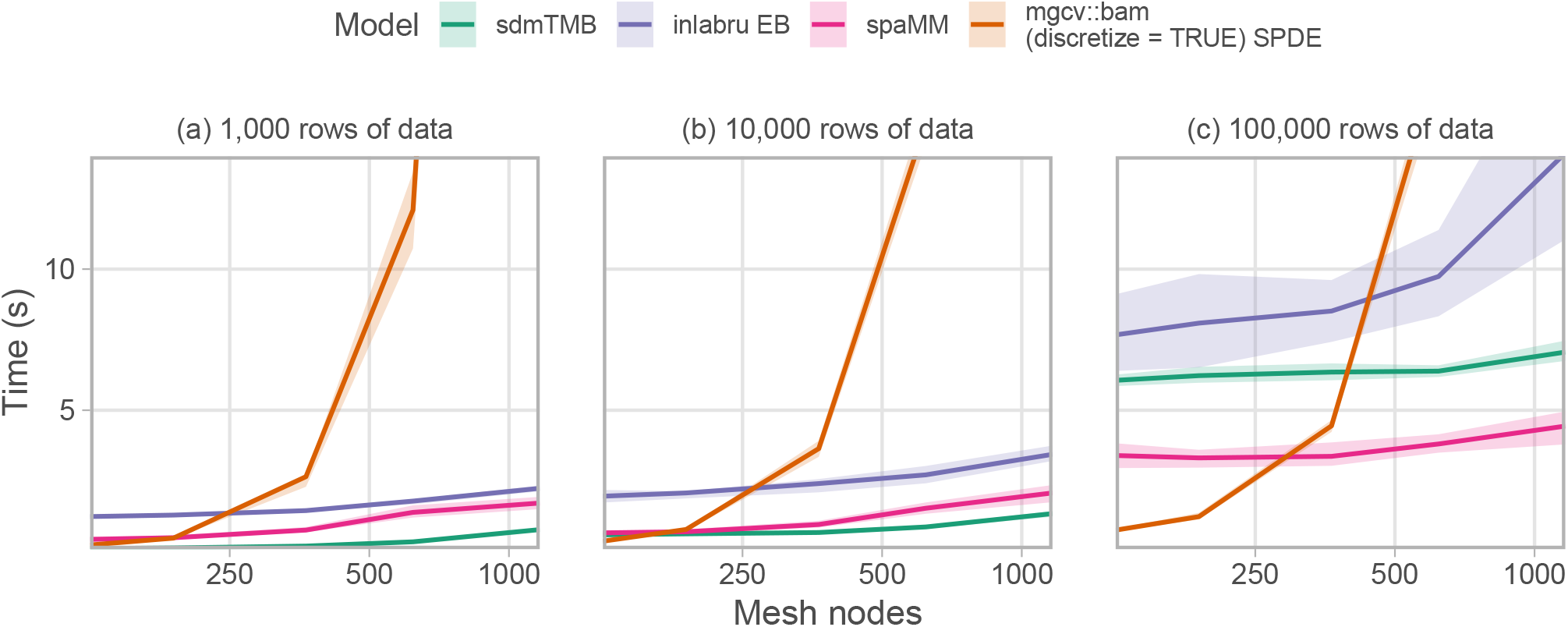
Comparison of time to fit an SPDE spatial random field model with an intercept, one fixed-effect predictor, Gaussian error, and a sequence of SPDE resolutions to three dataset sizes. Lines represent means and ribbons 90% quantiles across 50 random iterations. **VAST** and **tinyVAST** should be similar to **sdmTMB** and so are not shown. **inlabru** (version 2.10.1) used the empirical Bayes integration strategy and Gaussian approximation with `bru_max_iter = 1`, and the `like()` formulation. **mgcv** (version 1.9.1) used `bam(), method = ‘fREML’`, and discretized covariates (Miller *et al*. 2019). Note that spaMM (version 4.4.16) only fits spatial, not spatiotemporal, models. All platforms were restricted to one core and could be faster with parallel computation or optimized algebra libraries. See Figure 13 for a version with log-log axes.

**Figure 13:**
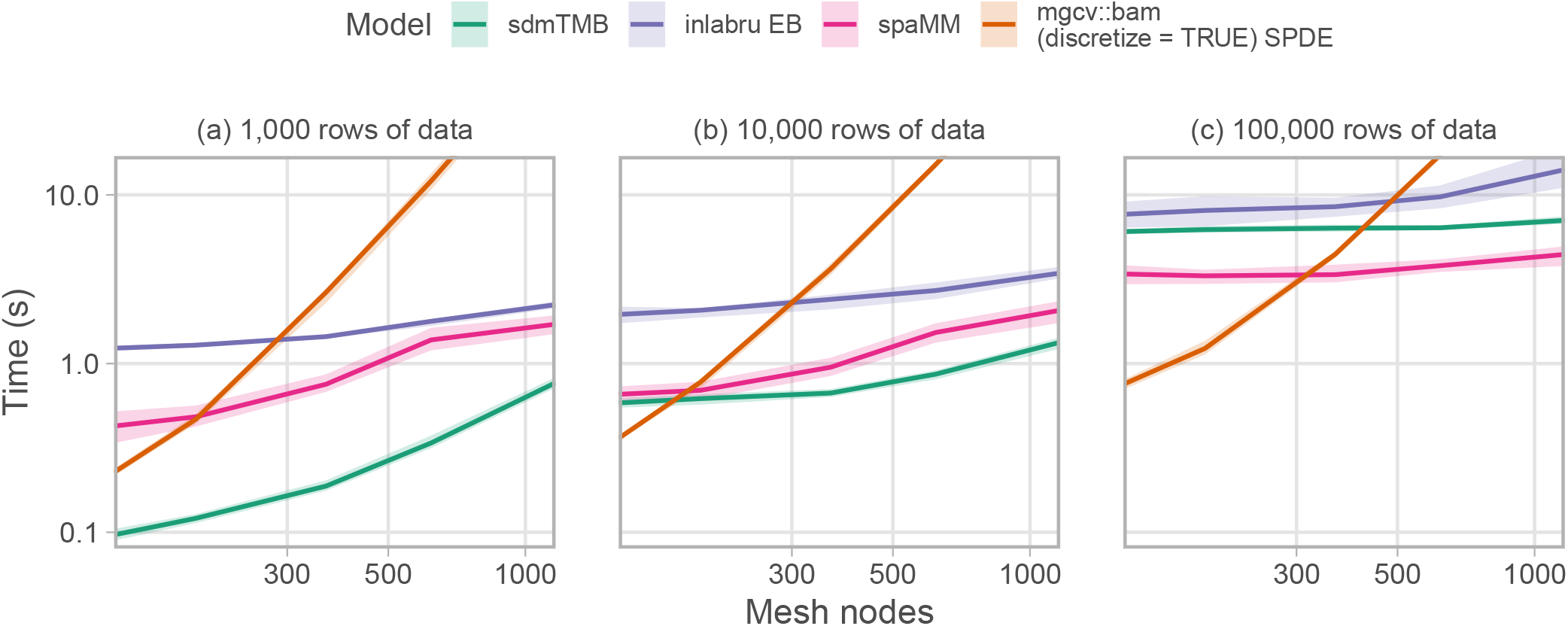
Same as Figure 12 but with both axes log distributed.

## 8. Discussion

How does one choose among the related packages mentioned in this paper to fit SPDE-based geostatistical GLMMs? Assuming a given package can fit the model of interest (Table 1), we suggest the primary differences are the user interface and speed. We think users famil-iar with stats::glm(), **lme4, mgcv**, or **glmmTMB** will find **sdmTMB** most approachable. Users familiar with R-INLA will find **inlabru** approachable. Users familiar with **mgcv** can adapt **mgcv** to fit similar models with custom code (Miller *et al*. 2019) and R-**INLA**/**inlabru** and **mgcv** are also general purpose modelling packages. VAST and **tinyVAST** are the sole options for fitting some multivariate models; alternatively, because these packages focus on multivariate delta models and fisheries applications, users fitting “simple” univariate spatial/spatiotemporal GLMMs will likely find **sdmTMB** more straightforward. Users looking for calculation, with uncertainty, of derived variables such as area-weighted population indexes, may **favour sdmTMB** or **VAST** (although such quantities can be calculated using other packages post hoc). Overall, **sdmTMB** unites functionality useful in many applied settings into a single package.

While our examples focused on applications to ecological data, the SPDE approach and the functionality of **sdmTMB** has applications in many other fields. Examples include spatial models of disease spread (Moraga, Dean, Inoue, Morawiecki, Noureen, and Wang 2021), spatial econometric models of quantities such as housing prices (Bivand, Gómez-Rubio, and Rue 2014), analyzing medical imaging data such as MRI scans (Parisa Naseri and Tabatabaei 2022), and geophysical models of seismic waves following earthquakes (Zhang, Czado, and Sigloch 2015). The **sdmTMB** model is further relevant to what is commonly referred to as spatial (Elhorst 2010; Lee and Yu 2010) and dynamic spatial panel data models (Elhorst 2012) in econometrics. These examples underscore the versatile nature of the SPDE approach and the potential uses of **sdmTMB** across various scientific and industrial sectors.

The GLMMs underpinning **sdmTMB** models are spatially explicit and derived from a mechanistic diffusion process—they estimate interpretable parameters of a spatial covariance function: parameters defining the magnitude of spatial variation and the rate of correlation decay with distance. In contrast, non-parametric approaches [e.g., **randomForest** (Liaw and Wiener 2002), **MaxEnt** (Phillips, Anderson, and Schapire 2006)] and most smooths in **mgcv** (Wood 2017) do not estimate spatial covariance functions. Approaches such as conditional autoregressive models (CAR) are applicable to areal data, where the spatial domain is discretized into a set of vertices or polygons. Areal data (data aggregated to polygon or grid level) may be analyzed using other spatial models, including CAR models (e.g., Ver Hoef, Peterson, Hooten, Hanks, and Fortin 2018); **sdmTMB** can also fit models with areal data if each polygon has an associated centroid. A benefit of the geostatistical approach over CAR or similar models is that the parameters describing spatial covariance can be more easily interpreted (Wall 2004).

Additional functionality in **sdmTMB** not already mentioned includes interpolating across missing time slices and forecasting, the barrier SPDE model (Bakka, Vanhatalo, Illian, Simpson, and Rue 2019), and time-varying spatiotemporal covariance parameters (Ward, Barnett, Anderson, Commander, and Essington 2022). There are several planned future additions to the **sdmTMB** model structure. A subset of features to be added include (1) the ability to specify observation-specific likelihoods to integrate different data types (e.g., Grüss and Thorson 2019), (2) inclusion of a dispersion formula similar to **glmmTMB**, (3) inclusion of random slopes to complement the existing random intercepts, and (4) integration with the **RTMB** (Kristensen 2023) package so that the model code base is written in R rather than C++.

## 9. Acknowledgements

**sdmTMB** would not be possible without the **TMB** (Kristensen *et al*. 2016) and R-INLA (Rue *et al*. 2009; Lindgren *et al*. 2011; Lindgren and Rue 2015) R packages. **sdmTMB** is heavily inspired by and in some places code has been adapted from both the **VAST** (Thorson 2019a) and **glmmTMB** (Brooks *et al*. 2017) R packages (as described in the DESCRIPTION and inst/COPYRIGHTS files). Smoother support was possible thanks to **mgcv** (Wood 2017). We thank the authors of all these packages. We thank S. Kotwicki, M. Lindmark, M. Martin, C.C. Monnahan, P.M. Regular, J. Watson, two anonymous reviewers, and the editor for helpful comments that substantially improved the manuscript. We thank the many individuals who have contributed to collecting the trawl survey data with Fisheries and Oceans Canada that was used in the Pacific Cod and Pacific Spiny Dogfish examples. Christmas Bird Count data for the Snowy Owl example were provided by National Audubon Society and through the generous efforts of Bird Studies Canada and countless volunteers across the Western Hemisphere.

## A. Speed testing related packages

Here, we describe the methods underlying the speed testing in Figure 12. We generated a mesh that was consistent across simulated data sets for a given mesh resolution (Figure 14). We did this by setting the `max.edge` argument, which controls the largest allowed triangle edge length. We tested values of `max.edge` of 0.06, 0.075, 0.1, 0.15, and 0.2. In Figure 12, we report on the x-axis the number of mesh nodes (“vertices”) that result from each of these meshes.

**Figure 14:**
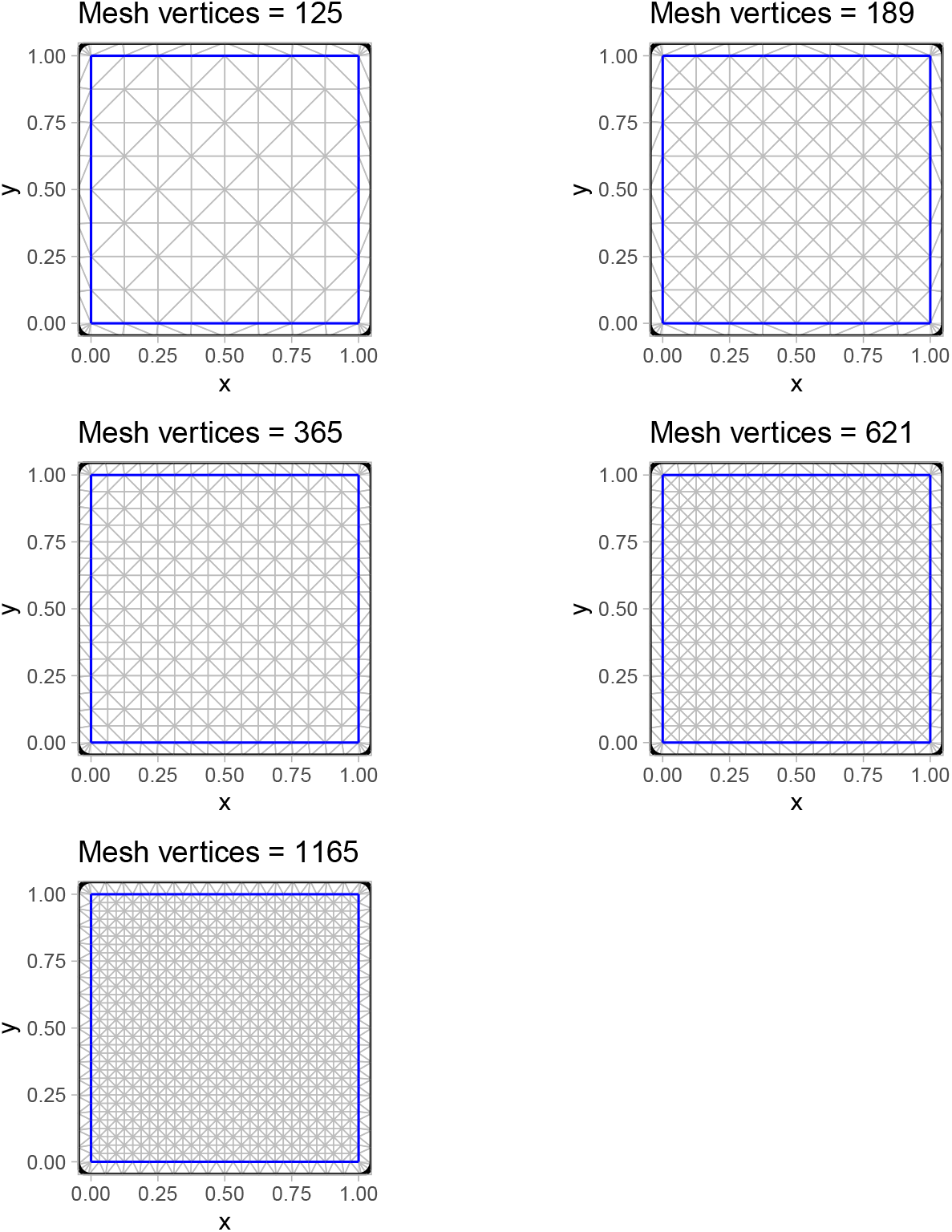
The meshes used in simulations from least to most vertices.

We simulated 1,000, 10,000, or 100,000 spatial observations with both x and y coordinates from uniform(0, 1) distributions (Figure 15). Each iteration generated unique coordinates, predictor data, Gaussian random field values, and observation error. The Gaussian random field was parameterized with a range of 0.5 and marginal standard deviation of 0.2. The model included an intercept with a value of 0.2 and a normal(0, 1) predictor with an associated coefficient of -0.4. The observation error was Gaussian with a standard deviation of 0.3.

**Figure 15:**
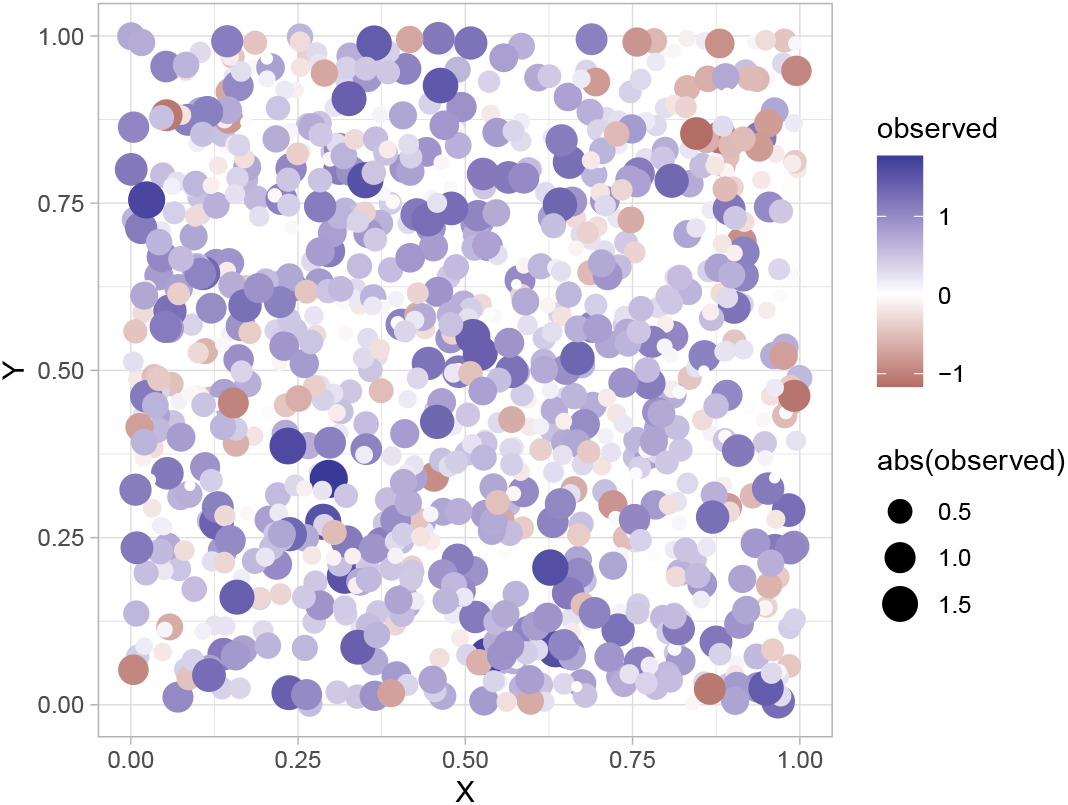
Example simulated dataset with a spatial random field.

We conducted 50 simulation iterations per model and mesh resolution and show timing results for the mean, 10%, and 90% quantile values. The models were fit on a 2023 16 inch M2 MacBook Pro with an Apple M2 Pro 10-core CPU and 32 GB of RAM in R 4.4.0 and the default **BLAS** library packaged with R(R Core Team 2024).

### A.1. Illustration of generating the INLA mesh

We will illustrate with a `max.edge` of 0.06:

~~~
*R*> *max_edge* <*- 0.06
R*> *loc_bnd* <*- matrix*(*c*(*0, 0, 1, 0, 1, 1, 0, 1*), *4, 2, byrow* = *TRUE*)
*R*> *segm_bnd* <*- INLA::inla.mesh.segment*(*loc_bnd*)
*R*> *mesh* <*- INLA::inla.mesh.2d*(
*+  boundary* = *segm_bnd*,
*+  max.edge* = *c*(*max_edge, 0.2*),
*+  offset* = *c*(*0.1, 0.05*)
*+*)
~~~

This mesh has 1165 (mesh$n) vertices.

### A.2. Illustration of simulating data

~~~
*R*> *set.seed*(*123*)
*R*> *n_obs* <*- 1000
R*> *predictor_dat* <*- data.frame*(
*+ X* = *runif*(*n_obs*), *Y* = *runif*(*n_obs*),
*+ a1* = *rnorm*(*n_obs*)
*+*)
*R*> *mesh_sdmTMB* <*- make_mesh*(*predictor_dat, xy_cols* = *c*(*“X”, “Y”*), *mesh* = *mesh*) *R*>
*R*> *sim_dat* <*- sdmTMB_simulate*(
*+ formula* = *∼ 1 + a1*,
*+ data* = *predictor_dat*,
*+ mesh* = *mesh_sdmTMB*,
*+ family* = *gaussian*(),
*+ range* = *0.5*,
*+ phi* = *0.3*,
*+ sigma_O* = *0.2*,
*+ B* = *c*(*0.2, -0.4*) *# B0* = *intercept, B1* = *a1 slope
+*)
~~~

### A.3. Example sdmTMB model fit

~~~
*R*> *fit_sdmTMB* <*- sdmTMB*(
*+ observed ∼ a1*,
*+ data* = *sim_dat*,
*+ mesh* = *mesh_sdmTMB*,
*+ family* = *gaussian*(),
*+ priors* = *sdmTMBpriors*(
*+ matern_s* = *pc_matern*(*range_gt* = *0.05, sigma_lt* = *2*)
*+*)
*+*)
~~~

### A.4. Example spaMM model fit

~~~
*R*> *spde* <*- INLA::inla.spde2.pcmatern*(
*+ mesh* = *mesh*,
*+ prior.range* = *c*(*0.05, 0.05*),
*+ prior.sigma* = *c*(*2, 0.05*)
*+*)
*R*>
*R*> *fit_spaMM* <*- fitme*(
*+ observed ∼ a1 + IMRF*(*1* | *X + Y, model* = *spde*),
*+ family* = *gaussian*(),
*+ data* = *sim_dat
+*)
~~~

### A.5. Example inlabru model fit

~~~
*R*> *dat_sp* <*- sp::SpatialPointsDataFrame*(
*+ cbind*(*sim_dat$X, sim_dat$Y*),
*+ proj4string* = *sp::CRS*(
*+  “+proj*=*aea +lat_0*=*45 +lon_0*=*-126 +lat_1*=*50 +lat_2*=*58.5 +x_0*=*1000000
+ + +y_0*=*0 +datum*=*NAD83 +units*=*km +no_defs”
+*), *data* = *sim_dat
+*)
*R*> *components* <*- observed ∼ -1 + Intercept*(*1*) *+ a1 +
+  spatrf*(*main* = *coordinates, model* = *spde*)
*R*> *like* <*- like*(*observed ∼ Intercept + a1 + spatrf*,
*+  family* = *“gaussian”, data* = *dat_sp
+*)
*R*> *fit_bru* <*- bru*(
*+ like*,
*+ components* = *components*,
*+ options* = *bru_options*(
*+  control.inla* = *list*(*int.strategy* = *“eb”, strategy* = *“gaussian”*),
*+  bru_max_iter* = *1, num.threads* = *“1:1”
+*)
*+*)
~~~

### A.6. Example mgcv model fit

First define smooth.construct.spde.smooth.`spec()` and Predict.matrix.spde.`smooth()` from the supplement of Miller *et al*. (2019), then:

~~~
*R*> *class*(*mesh*) <*- “inla.mesh”
R*> *fit_bam* <*- bam*(
*+ observed ∼ a1 + s*(*X, Y*,
*+  bs* = *“spde”, k* = *mesh$n*,
*+  xt* = *list*(*mesh* = *mesh*)
*+*),
*+ data* = *sim_dat*,
*+ family* = *gaussian*(),
*+ method* = *“fREML”*,
*+ control* = *gam.control*(*scalePenalty* = *FALSE*),
*+ discrete* = *TRUE
+*)
~~~

